# Gut Microbiome of Helminth Infected Indigenous Malaysians Is Context Dependent

**DOI:** 10.1101/2022.01.21.477162

**Authors:** Mian Zi Tee, Yi Xian Er, Alice V. Easton, Nan Jiun Yap, Ii Li Lee, Joseph Devlin, Ze Chen, Kee Seong Ng, Poorani Subramanian, Angelina Angelova, Shushan Sargsian, Romano Ngui, Christopher Chiong Meng Boey, Kek Heng Chua, Ken Cadwell, Yvonne Ai Lian Lim, P’ng Loke, Soo Ching Lee

**Affiliations:** Department of Biomedical Science, Faculty of Medicine, Universiti Malaya, Kuala Lumpur, Malaysia; Department of Parasitology, Faculty of Medicine, Universiti Malaya, Kuala Lumpur, Malaysia; Department of Microbiology, New York University Grossman School of Medicine, New York, NY, United States; Kulliyyah of Medicine and Health Sciences, University Islam Antarabangsa Sultan Abdul Halim Mu’adzam Shah, 09300 Kuala Ketil, Kedah; Department of Gastroenterology, Faculty of Medicine, Universiti Malaya, Kuala Lumpur, Malaysia; Bioinformatics and Computational Biosciences Branch, Office of Cyber Infrastructure and Computational Biology, National Institute of Allergy and Infectious Diseases, National Institutes of Health, Bethesda, MD, USA; Kimmel Center for Biology and Medicine at the Skirball Institute, New York University Grossman School of Medicine, New York, NY, United States; Department of Paediatrics, Faculty of Medicine, Universiti Malaya, Kuala Lumpur, Malaysia; Division of Gastroenterology, Department of Medicine, New York University Langone Health, New York, NY, United States; Type 2 Immunity Section, Laboratory of Parasitic Diseases, National Institute of Allergy and Infectious Diseases, National Institute of Health, Bethesda, MD, United States

**Keywords:** helminth, microbiome, metagenomic sequencing, indigenous population, albendazole

## Abstract

While microbiomes in industrialized societies are well characterized, indigenous populations with traditional lifestyles have microbiomes that are more akin to those of ancient humans. However, metagenomic data in these populations remains scarce and the association with soil-transmitted helminth infection status is unclear. Here, we sequenced 650 metagenomes of indigenous Malaysians from 5 villages with different prevalence of helminth infections. Individuals from villages with higher prevalence of helminth infections have more unmapped reads and greater microbial diversity. Microbial community diversity and composition were most strongly associated with different villages and the effects of helminth infection status on the microbiome varies by village. Longitudinal changes in the microbiome in response to albendazole anthelmintic treatment was observed in both helminth infected and uninfected individuals. Inference of bacterial population replication rates from origin of replication analysis identified specific replicating taxa associated with helminth infection. Our results indicated that helminth effects on the microbiota was highly dependent on context and effects of albendazole on the microbiota can be confounding for the interpretation of deworming studies. Furthermore, a substantial quantity of the microbiome remains undescribed and this large dataset from indigenous populations associated with helminth infections should facilitate characterization of the disappearing microbiome from developed industrialized societies.

## Introduction

Industrialization is associated with reduced diversity of the microbiome in the human population [1], which could influence a range of physiological processes including nutrition, metabolism, immunity, neurochemistry and drug metabolism [2]. Traditional indigenous populations have substantially greater microbial diversity than individuals living in industrialized societies. Nonetheless, our current knowledge of the human gut microbiome [3] is overrepresented by data available from industrialized countries and does not fully address the under-sampling of indigenous populations, which may contain the largest source of disappearing microbes that could be utilized for restoration of the ancestral human microbiome [4].

Throughout evolution, helminths have coexisted with the gut microbiota in their mutual host [5] and the reduced prevalence of helminth infections from modern practices could contribute to the “hygiene hypothesis” [6]. By co-evolution, helminths and microbiota are likely to interact and affect each other, thus having significant effects on the host. While the effect of helminth colonisation on the human gut microbiota has been increasingly studied, the results reported have been inconsistent. Some studies find that helminth colonization changes gut microbial diversity and composition and/or a shift in abundance of certain bacterial taxa [7-13], while others show no apparent changes in gut microbial profiles [14, 15]. These divergent conclusions could be attributed to different confounders from different geographical locations (e.g. Malaysia [7, 13], Indonesia [10], Liberia [10], Tanzania [12], Western Kenya [11] and Ecuador [14]), different prevalence of helminth species (e.g. *Trichuris* sp. [14], hookworm [15], *Ascaris* sp. [11], *Strongyloides* sp. [9] and *Schistosoma* spp. [16]), as well as different approaches taken (natural or experimental infection, types of sequencing method and analysis approaches). Additionally, the direct impact of anthelmintic treatment on the gut microbiome is unclear. While some studies found differences following deworming treatment [11, 13], others have found no impact of treatment on gut microbiota profiles [14]. Other studies that examined anthelmintic albendazole effects on the gut microbiota utilize primarily 16S rRNA sequencing [10, 11, 14, 17, 18]. Hence a large study incorporating metagenomic sequencing with helminth infection status, albendazole treatment and additional control groups may provide greater insights into these complex interactions.

Most studies above utilized 16S rRNA sequencing to characterize the gut microbiota. Shotgun metagenomic approaches enable higher taxonomic resolution at the species level, and can identify not only bacteria, but also archaea, fungi and viruses [19, 20]. However, the deficiency of reference databases for mapping metagenomic data can make it a challenge to profile uncharacterized microorganisms. Recently, an approach of assembling sequencing reads into contigs and binning them into putative genomes, known as metagenome assembled genomes (MAGs), has enabled retrieving genomes directly from samples without the need of culturing organisms [21, 22]. The Unified Human Gastrointestinal Genome (UHGG) established an integrated catalog of prokaryotic genomes containing 204,938 non-redundant genomes that represent 4,644 prokaryotic species [3] by combining recent studies with large scale assembly of MAGs from human microbiome data [22-24] as well as two culture-based studies that sequenced genomes from cultivated human gut bacteria [25, 26]. The Human Reference Gut Microbiome (HRGM) catalog expanded on UHGG to include under-represented Asian metagenomes from Korea, India and Japan [27] and added 780 new species from the newly assembled genomes [27]. However, Southeast-Asian countries remain under-represented.

In this study, we utilized shotgun metagenomics on 650 Malaysian stool samples to investigate helminth-gut microbiome interactions by both cross-sectional and longitudinal analyses. The large sample size allowed us to examine these interactions in 5 different villages from different locations and lifestyle. Examination of anthelmintic treated uninfected individuals in the longitudinal phase enabled assessment of albendazole effects on the gut microbiome. Metagenomic data enabled investigating the replication rates of individual bacterial species under different conditions. Since a substantial quantity of the microbiome remains undescribed, this large dataset from indigenous populations with traditional lifestyles from the under-represented South East Asian region provides new insights into helminth-gut microbiome interactions and more comprehensive metagenomic sequences for future human gut microbiome studies.

## Results

### Gut microbiome analysis of indigenous Malaysians and urban controls

To identify and characterize helminth-associated microbiome effects, this study consisted of a cross-sectional component that compares urban individuals (N=56) living in Kuala Lumpur (KL) with indigenous Orang Asli (OA) (N=351) from five different villages (Figures S1 & S2), as well as a longitudinal component to examine changes to the microbiome after anthelminthic (albendazole) treatment. A total of 650 fecal samples (including longitudinal samples) were analyzed by metagenomic sequencing generating 11,480,206,516 paired reads after quality control and contamination removal (Figure S3). We compared different OA villages, which have different prevalence of soil-transmitted helminth infections (Figures S1 & S2). In the longitudinal phase of the study, consented OA subjects were treated with 400mg albendazole for 3 consecutive days after collection of the first fecal sample. At 21- and 42-days following treatment, additional fecal samples were collected, however this phase of the study was disrupted by the COVID-19 pandemic, reducing the number of paired samples available. KL subjects were not treated with albendazole and provided only one sample. Questionnaire data were collected and analyzed for some of the study subjects (n=340).

When we first mapped the metagenomic sequences to RefSeq (i.e., bacteria, protozoa, fungi, viral, archaea genomes), we observed a very low percentage of mapped reads (median: 41.6%). However, when we mapped the sequences to databases that incorporate metagenome assembled genomes (i.e., HRGM [27] and UHGG [3], the percentage of sequencing reads mapped to HRGM (median: 91.5%) and UHGG (median: 87.9%) was much higher than RefSeq (Figure 1A). Additionally, the percentage of mapped reads to all three databases was higher in KL subjects than the OA population [HRGM: p= 2.6e^-11^; UHGG: p= 1.2e^-07^; RefSeq: p=2e^-07^] (Figure 1A), indicating that there are more unknown microbial genomes in the OA population.

**Fig. 1.**
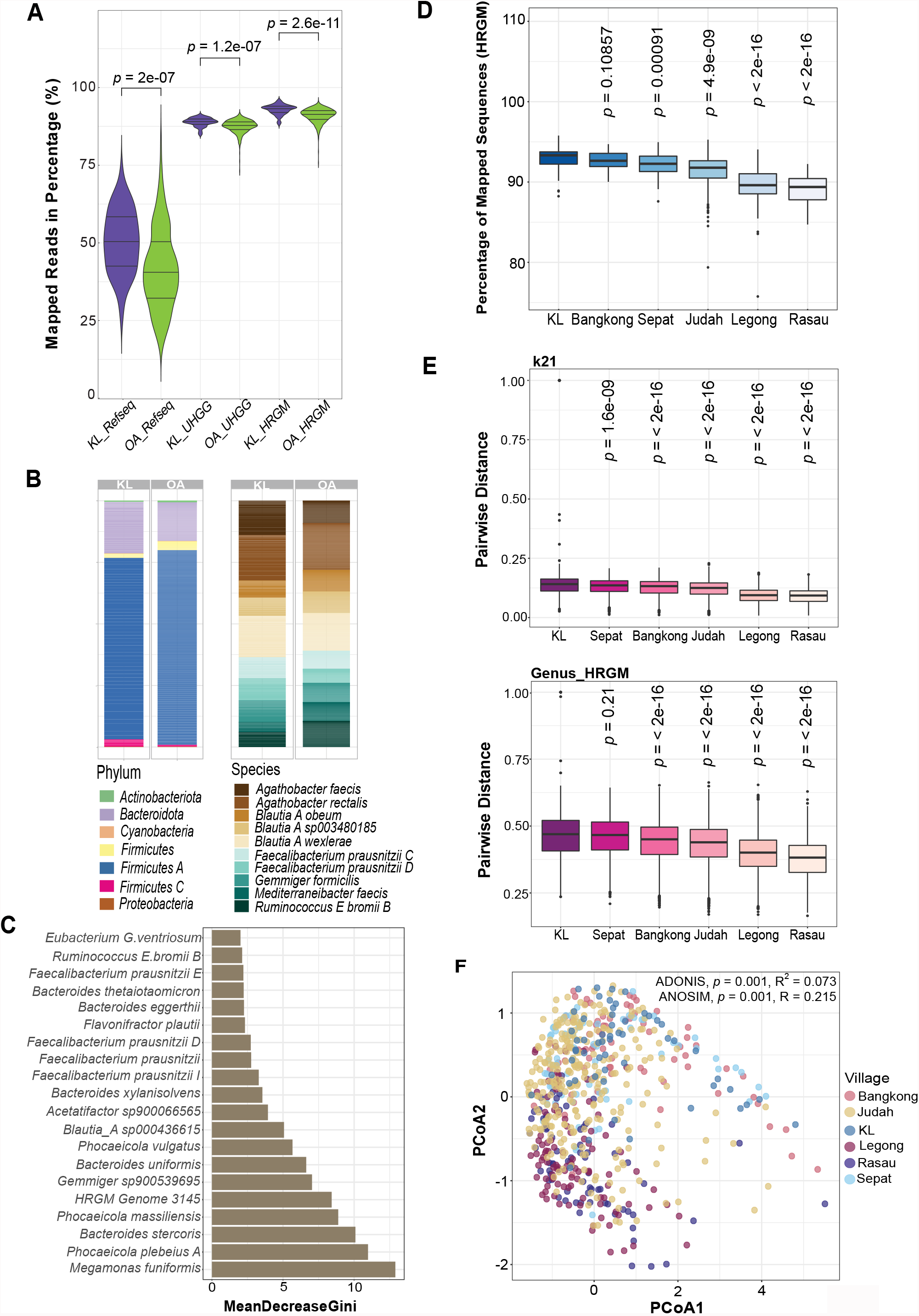
Variation in the gut microbiome of 650 Malaysians from Orang Asli (OA) villages and Kuala Lumpur (KL). **A** Violin plots illustrating the percentage of mapped reads with RefSeq (i.e., Bacteria, protozoa, fungi, viral, archaea), Unified Human Gastrointestinal Genome (UHGG) and Human Reference Gut Microbiome (HRGM) databases between OA (green) and KL (purple) samples. **B** Relative abundance of phyla from the 237 species of the core gut microbiota of OA and KL populations (Left). The relative abundance of the main species from Firmicute A (Right). **C** Bar plot of the top 20 species from the core gut microbiota that best predict the difference between OA and KL cohorts using a Random Forest classification model. **D** The percentage of mapped reads to the HRGM database for samples from different OA villages and KL. **E** Pairwise beta diversity comparisons of all villages to the KL cohort, assessed by Jaccard distance based on the distance of nucleotide k-mer sketches (k=21), at approximately genus level (Top) and genus-level classification (Bottom). **F** Principal Coordinates Analysis (PCoA) of Jaccard distance based on the gut metagenomic profiles in all samples, with individuals from different geographical locations denoted by specific color (ADONIS: p=0.001, R^2^=0.073; ANOSIM: p=0.001, R=0.215). The p-values for A, D, E is computed using Wilcoxon rank sum test.

Utilizing HRGM, we determined the core microbiota for the Malaysian population and found that 237 core bacterial species were 100% shared among the subjects (Figure 1B; Supplementary Figures S4A-E). The most abundant phylum was Firmicutes A, the majority of which were uncultured species [3] (Figure 1B). *Agathobacter rectalis, Balutia_A wexlerae* and *Agathobacter faecis* were the main species from Firmicutes A (Figure 1B). Using a cross-validated random forest model to identify core microbiota species driving the variation between OA vs KL subjects, we achieved a mean prediction accuracy of 98.05% at a kappa of 96.06% (out-of-bag error = 1.8%). *Megamonas funiformis, Phocaeicola pleneius A, Bacteroides stercoris, Phocaeicola massiliensis* and HRGM Genome 3145 were the top 5 predictors between OA and KL subjects (Figure 1C). Of these, HRGM Genome 3145, *Gemmiger sp900539695* and *Blautia A sp000436615* were more abundant in OA subjects, while *Megamonas funiformi, Phocaeicola pleneius A* and *Bacteroides stercoris* were more abundant in KL subjects (Supplementary Figure S5A & B). The bacterial species with the largest variation (cut-off 6.0 for the coefficient of variation) among the core gut microbiota is shown in Figure S6A.

The Orang Asli live in different geographical settings and have distinctive culture and lifestyle. We found that KL subjects have higher mapped reads than all OA villages (Figure 1D; Supplementary Figures S6B & C), and the percentage of mapped reads from both villages Rasau (p= 2e^-16^) and Legong (p= 2e^-16^) were markedly lower compared to KL (Figure 1D; Supplementary Figures S6B & C). Using a reference independent strategy to confirm our findings, we compared the beta-diversity (distributions of individuals pairwise distances in reference to KL) with k-mers 21 sketches at the genus level (Figure 1E), showing comparable results. In addition, we observed that Rasau and Legong had the highest beta-diversity and nucleotide dissimilarity compared to KL (Figure 1E). Moreover, comparison of bacterial communities across geographical locations using Jaccard distance revealed substantial differences between villages [ADONIS: p=0.001, R^2^=0.073; Analysis of similarity (ANOSIM): p=0.001, R=0.215] (Figure 1F, Supplementary Table S1). From the Principal Coordinate Analysis (PCoA) plot (Figure 1F), we observed clustering of the samples from Rasau and Legong. Conversely, the samples from Bangkong and Sepat were clustered together with KL while Judah exhibited a more dispersed distribution. Hence, OA subjects in Rasau and Legong were more similar in gut microbial composition, and were different from KL and other villages. Equivalent beta-diversity results were observed with other k-mers sketches (31 and 51) and at the species level (Figures S7A-G).

### Village dependent effects of helminth infection on the gut microbiome

We determined the infection intensity and the prevalence of intestinal helminth infection among the 351 OA subjects and found that *Trichuris* infection (61.8%, n=217) was the most predominant, followed by hookworm (20.8%, 73) and *Ascaris* (17.9%, 63) infections (Figure 2A). The distribution of age and gender of these subjects are shown in Figures S8A & B. The overall prevalence of helminth infection was 67.2% (n= 236) (Figure 2A) and infection intensity was summarized in Figure S8C. For beta diversity, based on PCoA, there were differences in gut microbiome between infected and uninfected individuals, however, statistically the effect size was small (ADONIS: p=0.001, R^2^=0.024; ANOSIM: p=0.001, R=0.145) (Figure 2B, Supplementary Table S1), which was also the case for Bray-curtis distance and NMDS ordination (Supplementary Figure S9A-C).

**Fig. 2.**
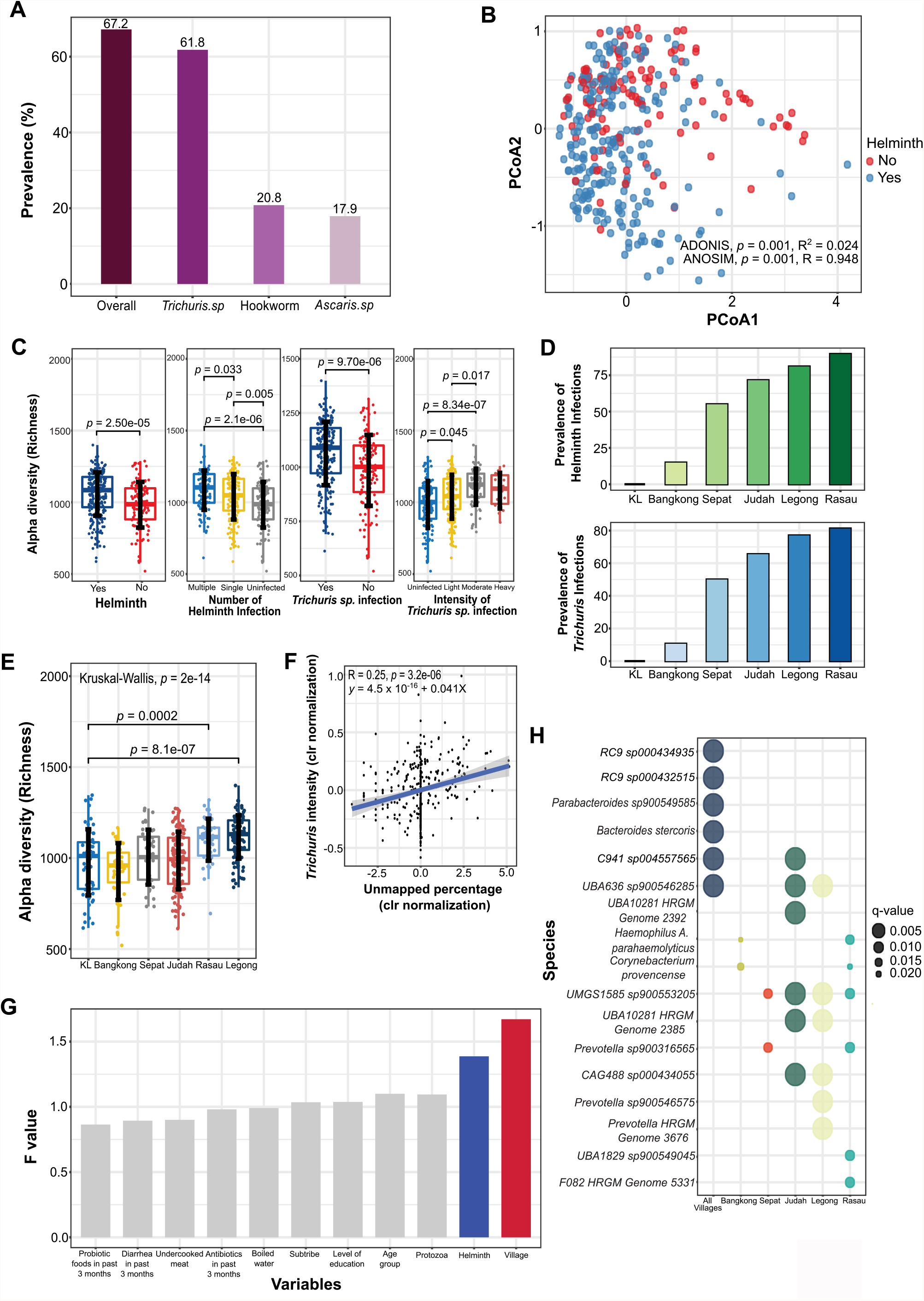
Effects of intestinal helminth infection status on gut microbial diversity and composition for the 407 Orang Asli individuals. **A** The prevalence of intestinal helminth infection in the OA cohort based on overall infection status, as well as specific intestinal helminth infection (i.e., Trichuriasis, ascariasis and hookworm infection). **B** Principal Coordinates Analysis (PCoA) of Jaccard distances based on gut microbiota profiles of the OA cohort. The individuals infected and uninfected with intestinal helminths are denoted by blue and red respectively (ADONIS: p=0.001, R^2^=0.024; ANOSIM: p=0.001, R=0.948). **C** Alpha diversity boxplot of species richness based on different status of intestinal helminth infection, number of intestinal helminth infection, *Trichuris* infection and intensity of *Trichuris* infection. Wilcoxon rank sum test is used for two independent variables while the Kruskal-Wallis test is used for more than two comparison groups. **D** The prevalence of intestinal helminth infection (Top) and *Trichuris* infection (Bottom) by different geographical locations. **E** Comparison of alpha diversity (species richness) between individuals from KL and specific OA villages. **F** Spearman correlation between the intensity of *Trichuris* infection and percentage of unmapped reads to the HRGM database (p=3.2e^-6^, R=0.25). The blue line represents the linear regression between intensity of *Trichuris* infection and percentage of unmapped reads. **G** Bar plot of the F statistic values from ADONIS analysis of variables that contribute to the gut microbiota composition. Colored bars indicate the variables that show significant effects on gut microbiota variation (p<0.05). **H** Bubble plot of bacterial species that are differentially abundant between *Trichuris* infected and uninfected individuals in all samples, as well as in specific village. The size of the bubble is negatively proportional to the p-value.

For alpha diversity, we observed higher species richness in the samples from infected subjects (p= 2.50e^-5^) (Figure 2C). Individuals infected with either single (p= 0.005) or multiple species of helminths (p= 0.033) had higher species richness (Figure 2C). *Trichuris* infected OA (p= 9.70e^-06^) had higher species richness than uninfected (Figure 2C), including those infected at light [eggs per gram (epg) < 999; p= 0.045] and moderate (epg < 9,999; p= 8.34e^-07^) intensities (Figure 2C). Other alpha diversity indices (i.e., Shannon and Simpson) are shown in Figures S10A-H and results for each village are shown in Figures S11A-E. The prevalence of helminth infection varied according to village and was highest in Rasau (89.6%, n= 43 of 48), followed by Legong (81.0%, 81 of 100), Judah (71.6%, 83 of 116), Sepat (55.0%, 22 of 40) and Bangkong (14.9%, 7 of 47) (Figure 2D). As *Trichuris* was the predominant helminth, the prevalence of *Trichuris* was similar for Rasau (81.3%, 39 of 48), Legong (77.0%, 77 of 100), Judah (65.6%, 76 of 116), Sepat (50.0%, 20 of 40) and Bangkong (10.6%, 5 of 47) (Figure 2D). There was no one infected with helminths in KL. The two villages with the highest prevalence, Rasau (p= 0.0002) and Legong (p= 8.1e^-07^), showed higher species richness compared to KL (Figure 2E). Also, we observed that species richness appeared to be greater when helminth infections in the villages were more prevalent, which was similar to the order of villages for unmapped reads shown in Figure 1D. To determine if *Trichuris* infection intensity was associated with unmapped reads, we performed a Spearman correlation test and found that the intensity of *Trichuris* infection was positively correlated (p=3.2e-06, R=0.25) with the percentage of unmapped reads to the HRGM database (Figure 2F). These results indicated that helminth infections were associated with under-representation in the catalog of bacterial genomes.

We next determined the relative contribution of village and helminth infection status on the gut microbiome in relation to other factors (e.g., if they had probiotic food, diarrhea, and antibiotics drug in the past 3 months, different age groups, subtribes and protozoa infection). ADONIS analysis indicated that only village (p=1.000e^-4^, F value= 1.672, R^2^= 0.025) and helminth status (p=0.028, F value= 1.387, R^2^= 0.010) had significant effects on the gut microbiome composition (Figure 2G). Since village has the largest effect size on gut microbiome composition, we next used MaAsLin2 [28] to identify bacterial taxa that were differentially abundant between *Trichuris* infected and uninfected individuals from specific villages. Importantly, we found that the bacterial species that were most differentially abundant between infected and uninfected subjects were unique to specific village (Figure 2H). For example, *Haemophilus_A*.*parahaemolyticus* and *Corynebacterium*.*provencense* were different in Bangkong and Rasau, whereas *Prevotella. sp900316565* was different in Sepat; C941.sp004557565 and UBA10281.HRGM_Genome_2392 in Judah; *Prevotella*.*sp900546575* and *Prevotella*.*HRGM_Genome_3676* in Legong; UBA1829.sp900549045 and F082.HRGM_Genome_5331 in Rasau (Figure 2H). Similar patterns of results were obtained with ANCOM-BC (Figure S12). These results indicated that helminth infections may have different effects on the gut microbiome in different villages.

### Dynamic changes to the gut microbiome after anthelminthic treatment

Longitudinal interventional approaches provide stronger assessment of cause-and-effect relationships. Fecal samples analyzed at Pre- and Post-anthelminthic treatment provided insights into the effects of deworming on the gut microbiome. Individual subjects were grouped into 4 categories [i.e., Full responders (n=43 paired; from 26-33,099 epg to 0 epg), Partial responders (n=23 paired; from 281-119,875 epg to 26-71,579 epg), Non-responders (n= 5 paired; from 204-1,097 epg to 281-1,632 epg) and Uninfected (n= 58 paired)], based on the *Trichuris* infection intensity before and after deworming (Figure 3A). While mixed infection was present in some individuals, hookworm and *Ascaris* infection were always cured after deworming (Supplementary Figure S13A).

**Fig. 3.**
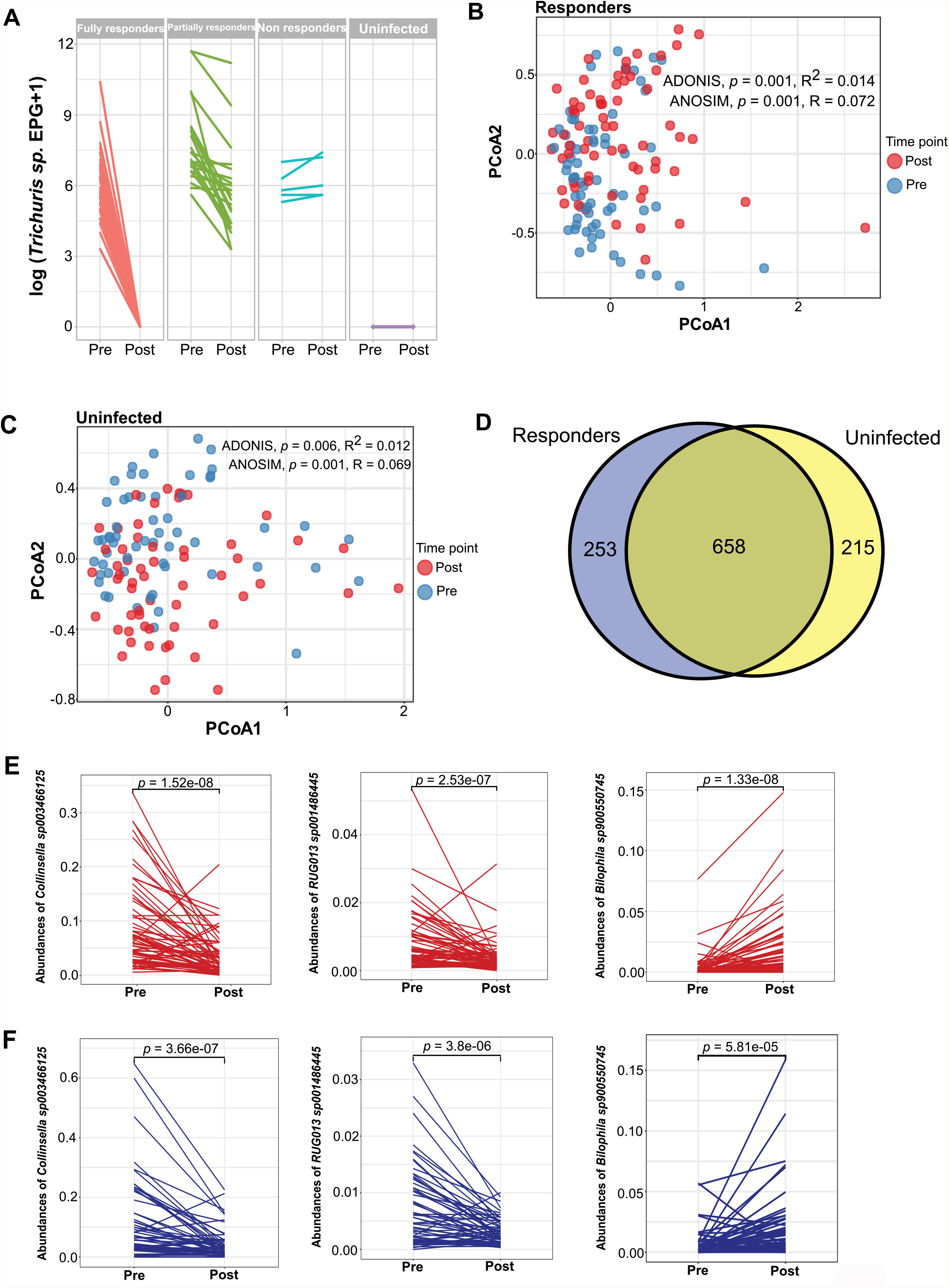
Dynamic changes to the gut microbiota of 129 Orang Asli after albendazole treatment. **A** Line plots show changes of the infection intensity of *Trichuris* pre and post response to anthelminthic drugs stratified by Full responders (n=43), Partial responders (n=23), Non-responders (n=5) and Uninfected individuals (n=58). **B** Principal Coordinates Analysis (PCoA) plot of Jaccard distances based on gut microbiota profiles of Responders (ADONIS: p=0.001, R^2^=0.014; ANOSIM: p=0.001, R=0.072), with Pre-anthelminthic treatment (blue) and Post-anthelminthic treatment (red). **C** Principal Coordinates Analysis (PCoA) plot of Jaccard distances based on gut microbiota profiles of Uninfected subjects (ADONIS: p=0.006, R^2^=0.012; ANOSIM: p=0.001, R=0.069) (Figure 3C, Supplementary Table S2), with Pre-anthelminthic treatment (blue) and Post-albendazole treatment (red). **D** Venn diagram depicting the number of shared and exclusive bacteria species that are found to be differentially abundant (Pre and Post) between Responders and Uninfected individuals. The blue area includes 253 bacteria that are altered only in Responders while the yellow and mixed color area indicates the 873 bacteria that are altered in Uninfected individuals. **E & F** Line plots showing changes to 3 of the the top differentially abundant bacterial species between Pre and Post treatment samples from **E** Responders and **F** Uninfected individuals, with p-values determined by the Wilcoxon signed-rank test. *Pre* Pre-anthelminthic treatment, *Post* Post-anthelminthic treatment.

First, we compared Pre and Post samples for Responders, which include both Full and Partial responders. PCoA based on Jaccard distances showed that there are differences in gut microbiota composition Pre and Post treatment, but the effect size was small (ADONIS: p=0.001, R^2^=0.014; ANOSIM: p=0.001, R=0.072) (Figure 3B, Supplementary Table S2). Since albendazole may have a direct effect on the microbiota, we next compared the gut microbiota profile Pre and Post treatment for helminth negative individuals. Similar to the Responders, PCoA based on Jaccard distances also indicated differences in gut microbiota composition between Pre and Post samples, with a small effect size (ADONIS: p=0.006, R^2^=0.012; ANOSIM: p=0.001, R=0.069) (Figure 3C, Supplementary Table S2). NMDS ordination, Bray-curtis distance matrix and beta-dispersion analysis showed similar results (Figures S14A-E and S15A-E, Table S2) and there were no significant changes to alpha diversity at Pre and Post treatment (i.e., Richness, Shannon, Simpson) (Supplementary Figures S13B and C).

Using MaAslin2 for differential abundance testing, we found changes of 911 bacterial species at Pre and Post treatment among Responders. However, there was substantial overlap with changes found in Pre and Post treatment samples for helminth negative individuals (658 species, 72.2%) (Figure 3D and Supplementary Figure S16), with only 253 taxa which were specific to the Responders. For example, in both Responders and uninfected individuals, the relative abundance of *Collinsella sp003466125* (p= 1.52e^-08^; p= 3.66e^-07^, respectively) and *RUG013*.*sp001486445* (p=2.53e^-07^; p= 3.80e^-06^) was reduced after deworming while the relative abundance of *Bilophila sp900550745* increased (p= 1.33e^-08^; p= 5.81e^-05^) (Figures 3E and F). Hence, albendazole may have a substantial effect on the microbiota that may be an important confounding factor for deworming studies.

In some individuals, we conducted a follow-up study 42-Days post-anthelminthic treatment. There were no differences in alpha diversity on Day-42 (Figure S17A) and although beta diversity analysis showed significant differences between three time points (i.e., Pre, 21-Day, and 42-Day) (Supplementary Figures S17B-C, Supplementary Table S3), these differences are driven by the pre-treatment samples (Supplementary Figure S17D). Therefore, the changes in the gut microbiome in both Responders and uninfected individuals after albendazole treatment remain stable by Day-42.

### Bacterial replication in the context of helminth infection

Actively replicating bacteria can be inferred based on calculating an index of replication from metagenomic sequencing for coverage trends resulting from bi-directional genome replication from a single origin of replication. We used the algorithm Growth Rate Index (GRiD) to estimate the growth rate of gut bacteria in relation to helminth infection status. We found that the growth rate of 61 bacterial species was associated with intestinal helminth infection (Figure 4A; Supplementary Table S4). Next, we conducted Spearman correlation analysis on *Trichuris* egg burden with the growth rate of these 61 bacterial species to identify the most highly correlated bacteria. From the 61 bacterial species, there were 11 bacterial species correlated to *Trichuris* egg burden (Figure 4B and Supplementary Figure S18A), with *Catenibacterium sp*. being positively correlated (p=2.83e^-04^, R=0.44) (Figure 4C) while *Bacteroides vulgatus* was negatively correlated (p=0.003, R=-0.39) (Supplementary Figure S18B). The growth rate of *Catenibacterium sp*. was notably higher in *Trichuris* infected individuals (p=1.65e^-16^). Conversely, the growth rate of *Bacteroides vulgatus* was lower in *Trichuris* infected individuals (p=2.65e^-16^) (Figure 4D).

**Fig. 4.**
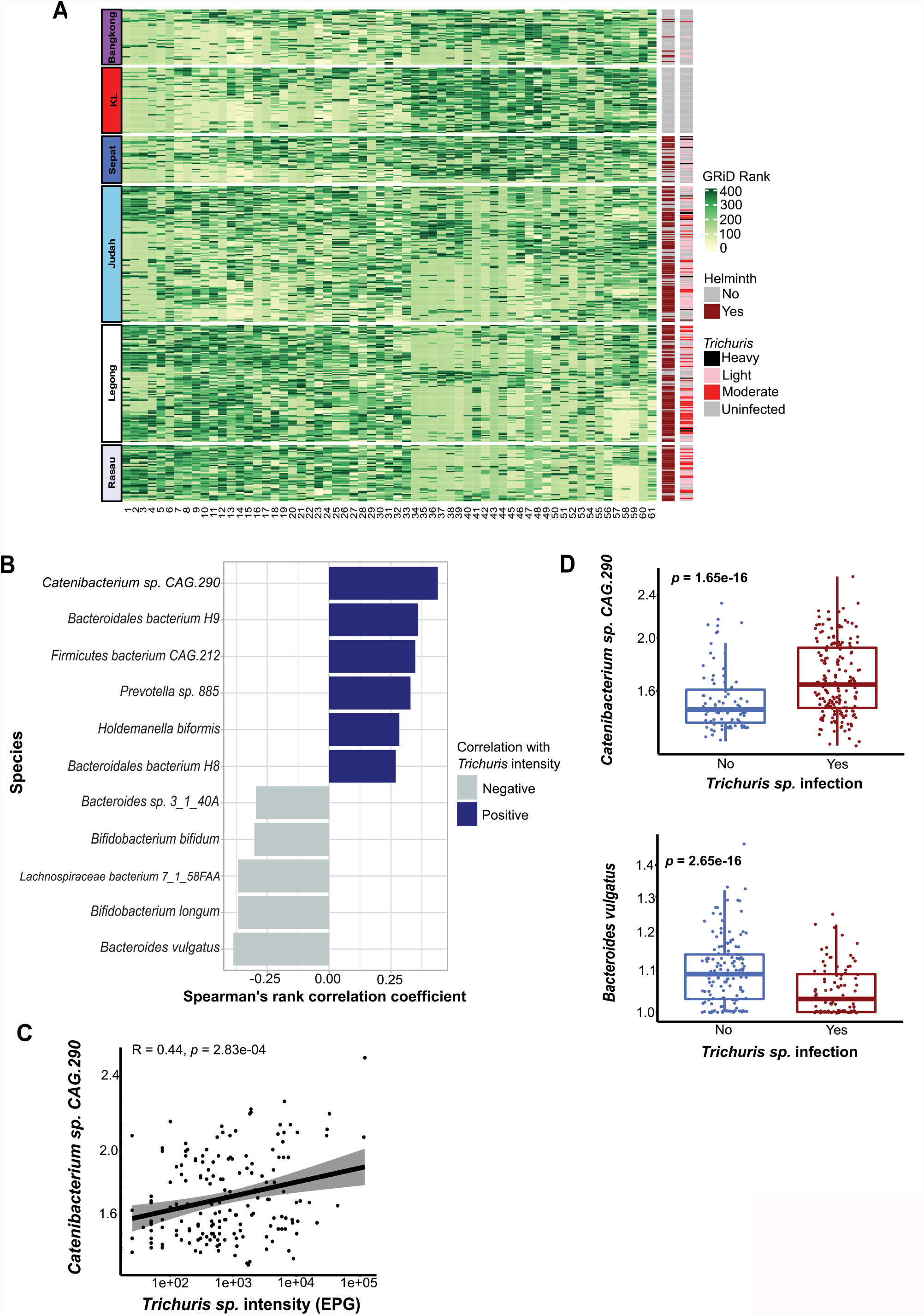
Gut bacterial replication in the context of intestinal helminth infection. **A** Heatmap of the Growth Rate Index (GRiD) score, which infers an index of replication for gut bacteria in relation to helminth infection status of individuals. Samples are shown in rows, by village, whereas the GRiD score of each bacterium are shown in columns. The first vertical side bar color codes the intestinal helminth infection status while the second side bar indicates the infection intensity of *Trichuris*. **B** GRiD score correlation between bacterial species with the infection intensity of *Trichuris*. The bar chart shows the Spearman’s rank correlation coefficient. Blue and grey colors represent the positive and negative correlations respectively. **C** Spearman’s rank correlation between *Catenibacterium sp*. growth rate and the infection intensity of *Trichuris* (p=2.83e^-4^, R=0.44). **D** Boxplots of GRiD score for *Catenibacterium sp*. and *Bacteroides vulgatus* in intestinal helminth infected and uninfected individuals. P-values are calculated by Wilcoxon rank sum test for cross-sectional comparison, Wilcoxon signed-rank test for longitudinal samples, with Benjamini-Hochberg correction.

For the longitudinal deworming component of the study, we observed that the growth rate of 12 bacterial species were different between Pre and Post treatment samples among the Responders. Among these bacterial species, half of them (i.e. *Bilophila wadsworthia, Lachnospiraceae bacterium_7_1_58FAA, Bacteroides vulgatus*, uncultured *Oscillibacter sp*., *Clostridium sp*.*CAG*.*81, Oscillibacter sp*.*57_20*) were also identified from the cross-sectional analysis (Supplementary Figure S18C). Spearman correlation analysis on *Trichuris* burden with the growth rate of these bacterial species demonstrated that uncultured *Oscillibacter sp*. (p=0.037, R= -0.29) and *Bacteroides vulgatus* (p=0.038, R= -0.24) were negatively correlated with *Trichuris* burden (Supplementary Figure S18D), indicating more replication in Responders after anthelminthic treatment (Supplementary Figure S18E). However, some of these relationships (*Firmicutes bacterium CAG*.*102, Lactobacillus rogosae, Clostridium sp. CAG*.*81* and *Bacteroides vulgatus*), were also observed in the helminth negative group (Supplementary Figure S18F). Hence, it could be difficult to disentangle the effects of helminth infection and direct effects of albendazole treatment on the dynamics of the microbiome.

## Discussion

In this study, we examined 650 stool metagenomes from a cohort of 351 indigenous Malaysians from 5 villages with different prevalence (14.9%-89.6%) of helminth infections, along with 56 urban citizens (uninfected) living in Kuala Lumpur city. To our knowledge, this is the largest study utilizing shotgun metagenomics to investigate the interactions between helminth infection and the human microbiome.

We found that mapping onto HRGM, which incorporates MAGs, improved taxonomic classification substantially, especially for the indigenous Orang Asli. We also found that the microbiota is dominated by Firmicutes A, which is represented by mostly uncultured bacteria, highlighting the under-representation of cultured bacteria from indigenous groups. This could be an important caveat for most of the previous studies on helminths and the gut microbiota, which were conducted using 16s rRNA sequencing [11-13, 17, 29] with the taxonomic classification based on mapping to the reference databases Greengenes, SILVA and Ribosomal Database Project (RDP). In a recent shotgun metagenomic study on 175 Cameroonian samples, the data was also mapped onto a reference database from NCBI [30] and the investigators noted that the classification of the relative abundance of bacteria did not correspond to data from 16S Greengenes classifications for the V4 region [30]. A different study also indicated that 16s rRNA gene sequencing only provided a portion of the gut microbiota profile compared to shotgun metagenomics [31]. Hence, we suggest assembling the Malaysian reference catalog to provide substantial benefit for future microbiome studies, especially from under-represented geographic regions and for rural and indigenous populations.

In this metagenomic study, we found that intestinal helminth infection status was associated with higher species richness, which was consistent with our previous findings and others conducted using 16s rRNA sequencing [10, 12, 13, 17, 30, 32]. However, we did not find a significant difference at Pre and Post deworming, which could be because of smaller sample size and could also be confounded by the effects of albendazole. Additionally, other studies have not observed an effect of helminths on microbial diversity [8, 11, 14, 17, 18, 33]. It is important to note that each study cohort has different prevalence rates for different helminth species, as well as distinct genetics and living conditions. This study has a larger sample size than our previous studies [13, 34] and enabled us to examine the interactions of helminth infection and the gut microbiome in different villages. Indeed, village has the largest effect size on gut microbiome variation, followed by helminth infection status. Notably, villages with higher helminth prevalence rates also have higher microbial diversity, but in different villages, helminth infection is associated with differential abundances of distinct bacterial taxa. It is important to note that the different villages represent diverse environments, practicing unique lifestyles and have different hygiene practices. Compared to other villages, Rasau and Legong villages (with higher helminth prevalence) are located near the forest with high exposure to the soil-environment, which may harbor more microbes [35] and mouse experiments showed that exposure to soil increases gut microbiota diversity [36]. From our questionnaire, a higher percentage of villagers from Rasau and Legong are plantation agricultural workers (Rasau: 30.4%; Legong: 14.8%; Others: < 6.7%), do not have a toilet (52.5%; 20.8%; <13.0%), and practice open defecation (46.5%; 32.0%; < 7.4%) more than other villages. As *Trichuris* eggs germinate in the soil this may increase exposure to *Trichuris*, as well as other microbes in the contaminated soil, resulting in higher microbial diversity in different settings.

We also found that deworming helminth negative individuals can influence the gut microbiome that overlaps substantially with changes in individuals responding to drug treatment by having reduced worm burdens. This indicates that albendazole may directly affect the microbiome, or that there are population effects that can influence uninfected people. There are four previous studies on albendazole [10, 11, 14, 17]. The first study conducted among Ecuador school children did not find any difference in bacterial composition among both *Trichuris* infected and uninfected groups after a combination of albendazole and ivermectin treatment [14]. In contrast, the second study in Indonesia found an increase of Actinobacteria and Bacteroidetes with albendazole treatment versus placebo in individuals that remained helminth-infected post-treatment, but not in uninfected individuals [17]. In addition, Rosa and coworkers also demonstrated that the gut bacterial composition was altered in a helminth-uninfected group in Indonesia after two years of albendazole treatment [10]. Another study in Kenya found significantly reduced Chao richness for uninfected individuals after deworming treatment, suggesting an effect of albendazole [11]. Albendazole is a prodrug that metabolizes rapidly to albendazole sulfoxide (the active anthelminthic compound) and albendazole sulphone (the inactive compound). Some bacterial species (*Enterobacter aerogenes* NCIM 2695, *Klebsiella aerogenes* NCIM 2258, *Pseudomonas aeruginosa* NCIM 2074 and *Streptomyces griseus* NCIM 2622) could be involved in metabolizing albendazole to albendazole sulfoxide and albendazole sulphone [37]. Albendazole can also be metabolized by the resident microbiota in gut rumens in sheep and cattle [38]. Hence, the gut microbiota could play a crucial role in metabolizing albendazole and influence drug bioavailability and efficacy on infected individuals. Why albendazole has lower efficacy against *Trichuris* infection than hookworm and *Ascaris* needs further investigation [39]. Future studies could apply metabolomics profiling to investigate metabolite differences between response groups to better understand the underlying mechanisms.

## Methods

### Study design and sample collection

This study consists of both cross-sectional and longitudinal phases. Cross-sectional comparisons were made on the Orang Asli cohort (OA) and between OA and urban cohorts (KL) living in the capital city of Malaysia, Kuala Lumpur. Within the Orang Asli community, we studied five Orang Asli villages: 1. Rasau village (Perak state); 2. Judah village (Selangor state); 3. Sepat village (Selangor state); 4. Bangkong village (Selangor state) and 5) Legong village (Kedah state). The locations of each village are displayed on a map using ArcGIS (version 10.7.1) together with other information including states, tribes and subtribes (Supplementary Figure S1). A total number of 351 samples were collected from Orang Asli subjects and 56 samples from KL subjects in this cross-sectional component (aged 4 years old and above) (Supplementary Figure S2).

For the longitudinal phase, Orang Asli subjects who provided consent were treated with 400mg albendazole for 3 consecutive days after the first stool sample collection. Stool samples were collected from the treated subjects at 21-days and 42-days following anthelminthic treatment. However, due to the restriction during the COVID-19 pandemic, only 4 Orang Asli villages were included in this phase, excluding Legong village. There was no follow-up for urban controls after the cross-sectional phase because they were not treated with albendazole. Sample selection for analysis was based on a complete set of paired stool samples [Pre (Pre-anthelminthic treatment) and Post (21-day post-anthelminthic treatment)] (n=129) and 3 time-points stool samples collection (Pre, 21-day and 42-day) (n=110). Four subject samples were removed from the longitudinal analysis due to incomplete data collection. Then, subjects were categorized into three groups for comparison: Responders, Non-responders and Uninfected, based on their infection status before and after the albendazole treatment. Responders (n= 66 paired samples) refer to individuals who were positive at baseline and became negative or showed reduction of infection intensity after deworming. Non-responders (n=5 paired samples) refer to individuals who were positive at baseline and showed increment or maintain of egg counts after deworming. Uninfected (n=8 paired samples) refer to negative individuals before and after the treatment. Non-responders were not be included in the gut metagenome analysis due to insufficient sample size. The details number of samples collected at each time points were shown in Supplementary Figure S2.

### Fecal sample preparation and analysis

All the stool samples collected were divided into two portions: (i) preserved in 2.5% potassium dichromate and stored at 4°C for intestinal helminth infection screening and (ii) aliquoted in 1.5ml cryovial tube, frozen immediately in dry ice and kept at -80°C for shotgun metagenomic analysis (Supplementary Figure S19). To detect and quantify helminth infections, Kato-Katz was performed. A thick smear was prepared from the fresh stool according to the manufacturer’s instructions (Kato-Katz kit, Mahidol University, Thailand) [40]. Infection intensity was stratified into light, moderate or heavy according to WHO cut-offs [41]. Formalin ether sedimentation was performed according to [42]. Stool samples were considered positive if any soil-transmitted helminths were detected from any of these two methods. DNA was extracted from stool samples using Qiagen DNeasy PowerSoil Pro kit (Qiagen, Hilden, Germany). DNA library were prepared using Illumina TruSeq DNA Nano Library kit (Illumina, United States). Paired end metagenomic sequencing was performed on the NovaSeq6000 S4 platform to generate an average of 20 million paired end reads per sample (range 13-35 million paired end reads), with a read length of 150bp and insert size of 350bp.

### Sequencing analysis pipeline

The overall bioinformatic analysis workflow from pre-processing to downstream analysis is shown in Supplementary Figure S3. In brief, the whole process of quality filtering and trimming of the raw sequence reads was performed by using KneadData (version 0.7.4) integrated with Trimmomatic [42], Bowtie [43] and FastQC [44] tools. Sequence reads were trimmed by using Trimmomatic with default settings, based on a sliding window trimming approach (SLIDINGWINDOW:4:20) when average base Phred quality score over four reads dropped below 33 (PHRED 33). Next, sequence reads were mapped against human reference genome (hg37) using Bowtie2 with default parameters (very-sensitive end-to-end alignment) to remove human host genome. The filtered reads were then used for the following analysis. Additionally, FastQC was used to perform quality checks on the raw metagenomic reads before pre-processing and after pre-processing to ensure of high-quality metagenomic reads for downstream analysis.

For taxonomic classification, Kraken2 (version 2.1.0) [45], a k-mer matching algorithm classifier was used for assigning taxonomic labels to the trimmed reads. The trimmed reads were mapped using Kraken2 against a standard database (1) RefSeq database (bacterial, protozoa, fungi, viral and archaeal) and two MAGs integrated databases (2) Human Reference Gut Microbiome (HRGM) database, with 232,098 reference genomes [27], (3) The Unified Human Gastrointestinal Genome (UHGG) database, with 204,938 reference genomes [24]. After taxonomic classification by Kraken2, Bayesian Re-estimation of Abundance with KrakEN (Bracken2) (version 2.6.0) [46] was used to compute the relative abundance of bacterial for each taxa (from phylum to species level) using a read length parameter of 150. The mapped reads of the OA and KL cohorts were then plotted into a violin plot using ggplot2 package [47] to access which databases provide better taxonomic classification. The distribution of the mapped reads was determined using the Shapiro test from the rstatix package [48]. Then, the difference between the mapped reads of KL was determined using the Wilcoxon rank sum test from ggplot2 package [47]. The data generated from Bracken2 were exported in the form of BOIM (Biological Observation Matrix) table and analyzed using R programming language (version 4.0.5, R Studio, Inc., Boston, MA, USA). The BIOM table was imported and filtered using the phyloseq package [49]. Only those taxa with a minimum abundance of 20% across all the samples and a minimum coefficient of variation of 3.0 were included in the following analysis (Supplementary Figure S20). In general, ggplot2 [47] and ggpubr package [50] was used to create visualization plots.

Sourmash (version 4.0.0) [51] was used to compute K-mer sketches. To discard erroneous kmers, the low abundance of k-mers were trimmed using “trim-low-abun” from khmer project, with a k-mer abundance cut-off of 3.0 and trimming coverage of 18. Signatures were generated for each sample using “sourmash compute” with a compression ratio of 10,000 (–scaled 10,000) and k-mer lengths of 21, 31 and 51 (-k21, -k31, -k51). A signature output was generated for Jaccard distance comparisons. Before the k-mer comparison, “sourmash index” was used to create a Sequence Bloom Tree (SBT) database from a collection of signatures. Lastly, “sourmash compare” was used with default settings to compare the signatures at each length of k.

The core microbiota was determined by including taxa present across all samples (i.e, abundance of 100% across all the samples). Random Forest (randomForest package) was used to identify microbiome taxa predictive of OA and KL [52] groups. Because of imbalance in the number of OA (n=594) and KL (n=56) samples, a “SMOTEd” (consist of 280 OA and 336 KL) data set was generated using the package DMwR [53] to counter this issue [54]. After that, the Random Forest model was built based on this “SMOTEd” data set, and tuned with the methods described by Brwonlee (2016) [55], followed by the significant testing using the methods described by [56].

Alpha diversity, in terms of species richness [57], Shannon [58] and Simpson index [59], were analyzed using the microbiomeSeq package [60]. The Wilcoxon rank sum test [61] was performed to compare groups statistically in the cross-sectional study (i.e., Helminth infected vs Uninfected and OA villages versus the KL) whereas Wilcoxon signed-rank test was used for paired samples in longitudinal study (i.e. Pre vs Post for both Responders and Uninfected). The Spearman’s rank correlation test [62] between the intensity of the *Trichuris* infection and the microbiome’s species richness was conducted after the data were normalized via centred log ratio transformation using the compositions package [63].

Beta diversity analysis was performed on both the Jaccard and Bray-Curtis dissimilarity matrix calculated from the taxon abundance data standardized using Hellinger. Differences in beta diversity between groups (i.e., different OA villages and different helminth infection status) or between different time points (Pre vs Post) were displayed with principal coordinates analysis (PCoA) plots and non-multi-dimensional scaling (NMDS) plots. The comparison on pair-wise distance of the samples between OA villages and KL was conducted using Wilcoxon rank sum test. This same analysis was also applied to the output generated from K-mers sketches.

Permutational multivariate analysis of variance (PERMANOVA) under ADONIS function [64] from the vegan package was conducted with 10,000 permutations on both the Jaccard and Bray-Curtis dissimilarity matrix. This analysis was first performed on specific variables of interest (i.e., different geographical locations, helminth status and Pre vs Post). ADONIS was used to assess the effect of multiple variables on the gut microbial composition (e.g., if they had probiotic food, diarrhea, and antibiotics drug in the past 3 months, different age groups, subtribes and protozoa infections), as well as Analysis of similarity (ANOSIM) [65]. To test for multivariate dispersions among groups, the permutation multivariate analysis of dispersion (PERMDISP) [64] was performed via the betadisper function and Tukey test under the Vegan package [66].

Differential abundance analysis was performed using Multivariate Association with Linear Models 2 (MaAsLin2) [28] for both cross-sectional (i.e., *Trichuris* infected vs non-infected for all villages and specific village) and longitudinal data (i.e., Pre vs Post for Responders and Uninfected). Analysis of composition Microbiomes with Bias Correction (ANCOM-BC) [67] was also conducted to validate the output generated from MaAsLin2, for cross-sectional data only. The Wilcoxon rank sum test and Wilcoxon signed-rank test from the rstatix package [48] were used to determine the p-value between groups for specific taxa.

Growth rate index (GRiD) (version 1.3) was used to evaluate the growth rate of microbial species in metagenomic samples [68]. Samples were mapped to a GRiD database (ftp://ftp.jax.org/ohlab/GRiD_environ_specific_database/stool_microbes.tar.gz), a stool-specific database created based on microbes mostly found in stool. GRiD score > 1.02 indicates bacteria are in growth phase whereas GRiD score < 1.02 indicates that bacteria are in stationary or lag phase. Bacterial species with GRiD score <1.02 in more than 75% of samples were filtered from further analysis, and then filtered using a cut-off of 2.0 of coefficient of variation (Supplementary Figure S21).

Cross-sectional comparison of bacterial species growth rate between helminth infected and uninfected individuals was computed using Wilcoxon rank sum test and corrected using Benjamini-Hochberg with false discovery rate (fdr) of 5%. Bacterial taxa that significantly differed (*p*<0.05) in GRiD score between helminth infected and uninfected individuals were displayed by heatmap. The top 20 taxa with the lowest p values were identified for Spearman’s rank correlation test with *Trichuris* infection intensity. This analysis was also applied to the longitudinal study to investigate the microbial growth rate between Pre and Post anthelminthic treatment among Responders and Uninfected individuals.

## Key Resources Table

List of the bioinformatic tools and R packages used are displayed in the table below:

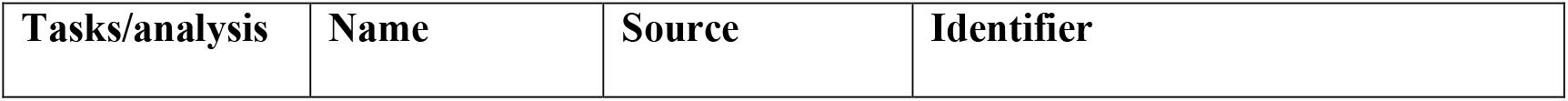
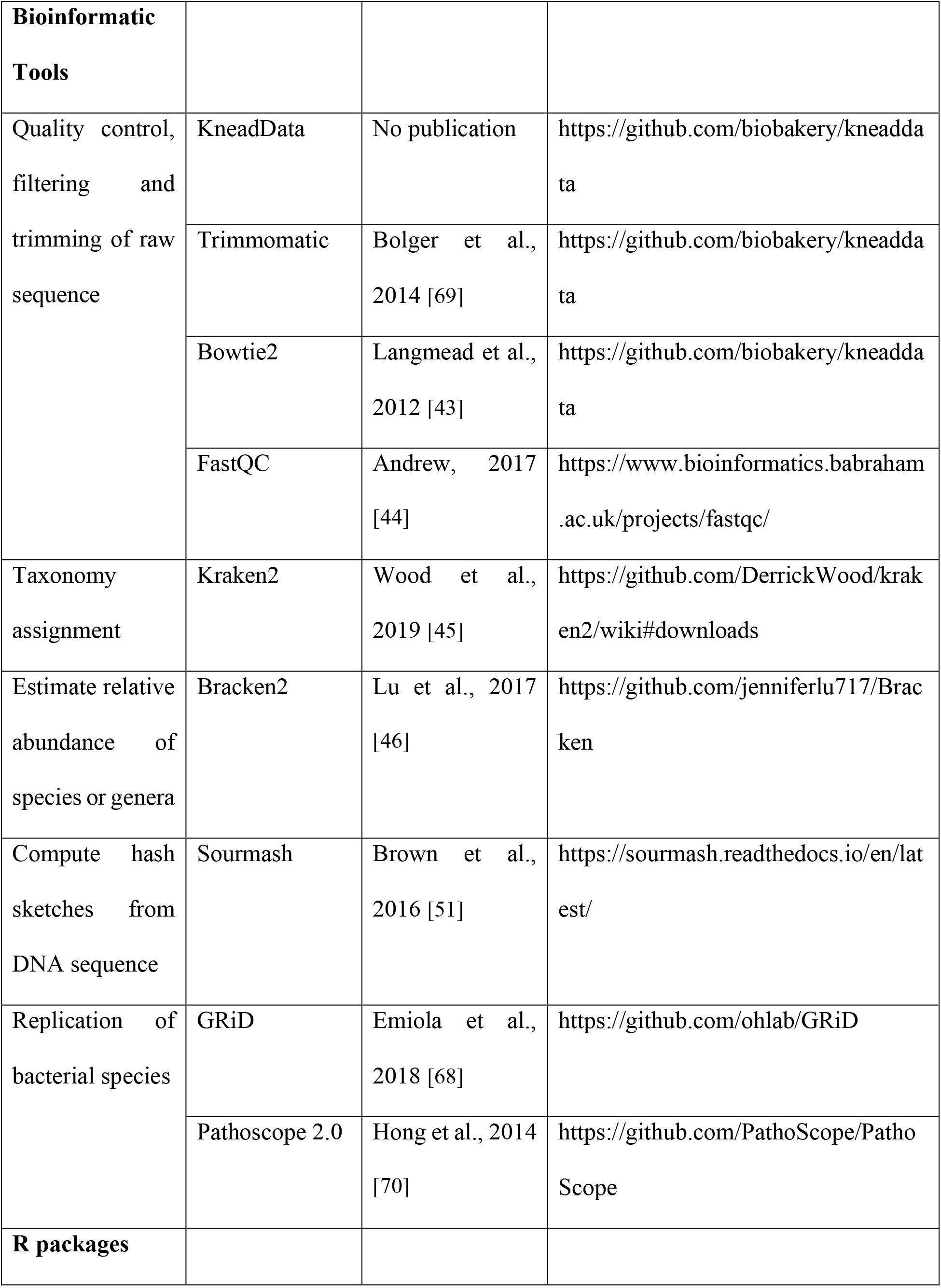
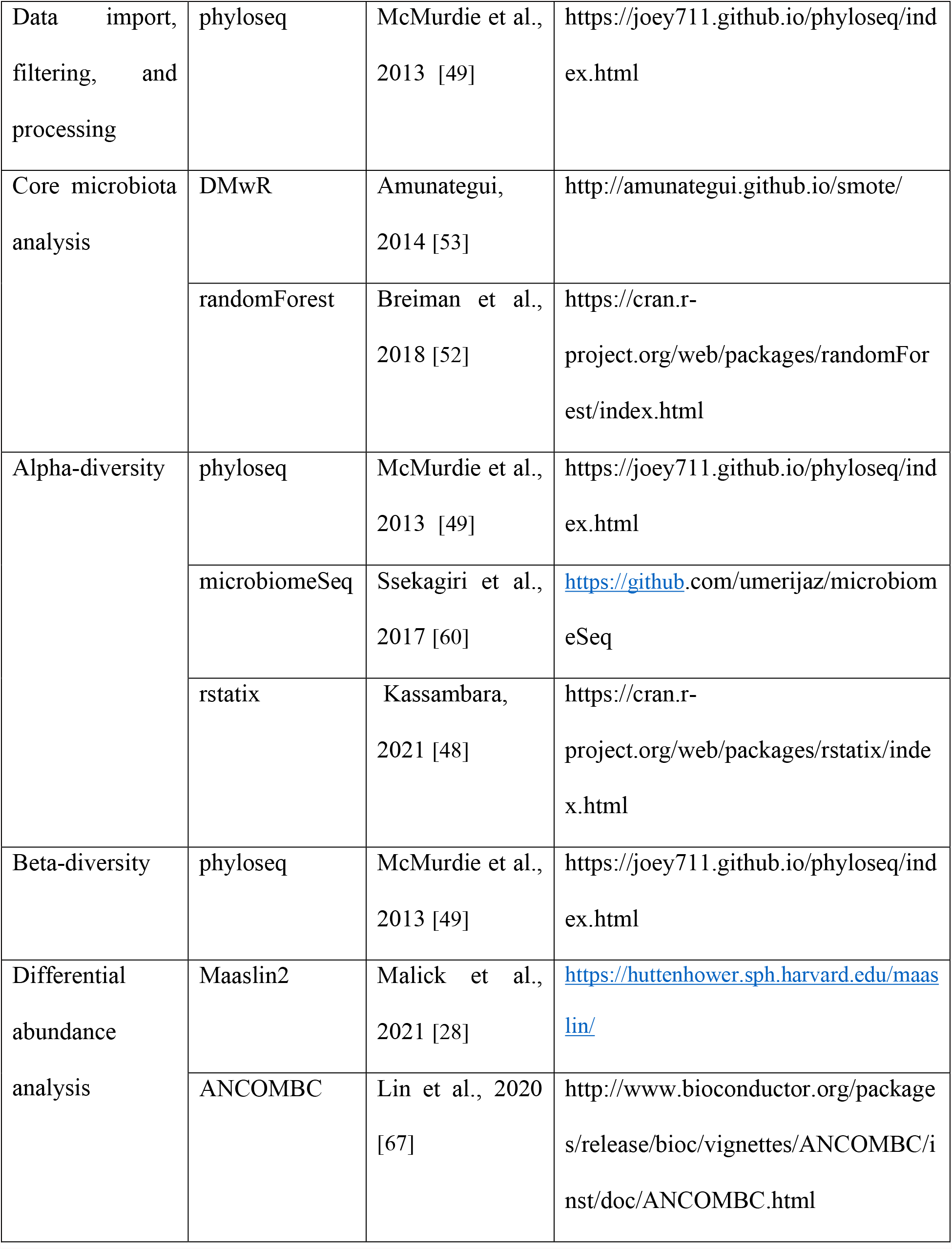
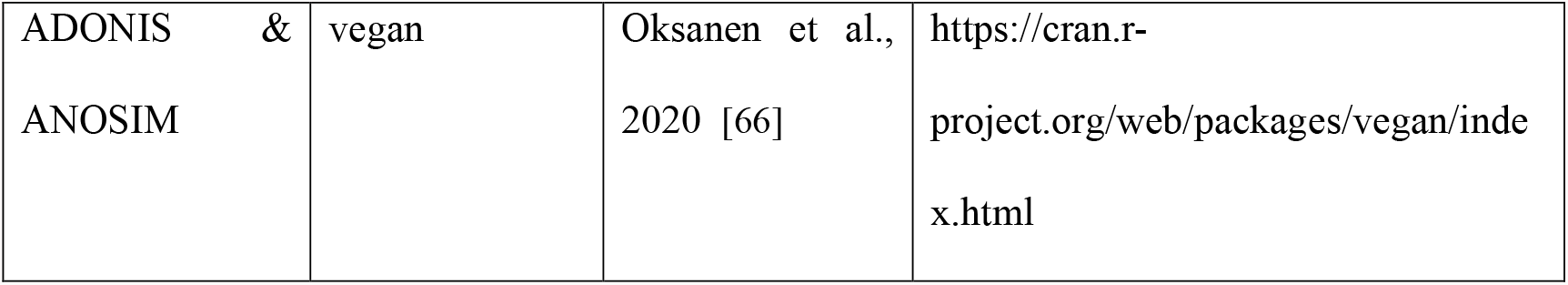

## Ethics approval and consent to participate

This study was approved by the Medical Ethics Committee of Universiti Malaya Medical Centre (UMMC) (Reference No.: 2017925-5593), National Medical Research Register (NMRR), Ministry of Health, Malaysia (Reference No.: NMRR-17-3055-37252), New York University (NYU) Institutional Review Board (IRB) (Reference No: i17-01068), the Department of Orang Asli Development (JAKOA) [Reference No.: JAKOA/pp.30.052Jld13 (12) & JAKOA/pp.30.052Jld14 (47)] and the chiefs of respective villages (Tok Batin) before enrolling the indigenous community in this study. An oral briefing using Malay language (the national language for Malaysia) on the purpose and the procedure of this study were explained to all the participants by the investigator. Written consent was attained from all adult participants aged 18 and above. For children under 18 years old, written parental consent was obtained from their respective parents or guardian and assent form was obtained from children aged 7 to 17 years old. Study exclusion criteria consisted of pregnant women, breastfeeding mothers and presence or perceived presence of any clinically significant disease.

## Availability of data and material

Raw data of gut metagenome has been deposited on the NCBI Sequence Read Archive (SRA) with the BioProject No. PRJNA797994 and BioSample accession No. SAMN25042866-25043515.

## Acknowledgements

We gratefully acknowledge JAKOA at the Ministry of Rural and Regional Development, Kuala Lumpur and the Village Chiefs for granting us permission to conduct this study and their cooperation during the whole course of study. We also thank all the participants for their voluntary participation and commitment in this study. Special thanks extended to the postgraduate and undergraduate volunteers for their valuable assistance during the fieldtrips. Further funding was provided by Faculty Scholar grant from Kenneth Rainin Foundation (K.C.) and Judith & Stewart Colton Center of Autoimmunity (K.C.). This work utilized the computational resources of the NIAID/NIH high performance computing (HPC) LOCUS cluster (https://locus.niaid.nih.gov/userportal/login.php?redirect=userportal%2Findex.php).

## Author contributions

PL, YALL & KC conceived and designed the study; SCL, MZT, YXE & ILL organized and coordinated field work; SCL, MZT, YXE, NJY & AVE carried out and oversaw the collection of samples from Malaysia; KSN provided clinical advice during sample collections; MZT & YXE performed the experiments; SCL, MZT & YXE analyzed and interpretation of all experimental data; JD, ZC, PS & AA advised and assisted in gut metagenome analysis; RN provided advice on mathematical analysis; CCMB, KHC & SS reviewed and edited the paper; PL, SCL, MZT & YXE wrote the paper with input from all authors.

## References

1. Cho I, Blaser MJ. The human microbiome: at the interface of health and disease. Nat Rev Genet. 2012;13(4):260–70.

2. Lynch SV, Pedersen O. The Human Intestinal Microbiome in Health and Disease. N Engl J Med. 2016;375(24):2369–79.

3. Almeida A, Nayfach S, Boland M, Strozzi F, Beracochea M, Shi ZJ, et al. A unified catalog of 204,938 reference genomes from the human gut microbiome. Nat Biotechnol. 2021;39(1):105–114.

4. Blaser MJ. The Past and Future Biology of the Human Microbiome in an Age of Extinctions. Cell. 2018;172(6):1173–77.

5. Loke P, Lim YA. Helminths and the microbiota: parts of the hygiene hypothesis. Parasite Immunol. 2015;37(6):314–323.

6. Blaser MJ, Falkow S. What are the consequences of the disappearing human microbiota? Nat Rev Microbiol. 2009;7(12):887–94.

7. Ramanan D, Bowcutt R, Lee SC, Tang MS, Kurtz ZD, Ding Y, et al. Helminth infection promotes colonization resistance via type 2 immunity. Science. 2016;352(6285):608–12.

8. Jenkins TP, Rathnayaka Y, Perera PK, Peachey LE, Nolan MJ, Krause L, et al. Infections by human gastrointestinal helminths are associated with changes in faecal microbiota diversity and composition. PLoS One. 2017;12(9):e0184719.

9. Jenkins TP, Formenti F, Castro C, Piubelli C, Perandin F, Buonfrate D, et al. A comprehensive analysis of the faecal microbiome and metabolome of Strongyloides stercoralis infected volunteers from a non-endemic area. Sci Rep. 2018;8(1):15651.

10. Rosa BA, Supali T, Gankpala L, Djuardi Y, Sartono E, Zhou Y, et al. Differential human gut microbiome assemblages during soil-transmitted helminth infections in Indonesia and Liberia. Microbiome. 2018;6(1):33.

11. Easton AV, Quiñones M, Vujkovic-Cvijin I, Oliveira RG, Kepha S, Odiere MR, et al. The Impact of Anthelmintic Treatment on Human Gut Microbiota Based on Cross-Sectional and Pre-and Postdeworming Comparisons in Western Kenya. mBio. 2019;10(2):e00519–19.

12. Chen H, Mozzicafreddo M, Pierella E, Carletti V, Piersanti A, Ali SM, et al. Dissection of the gut microbiota in mothers and children with chronic Trichuris trichiura infection in Pemba Island, Tanzania. Parasit Vectors. 2021;14(1):62.

13. Lee SC, Tang MS, Lim YA, Choy SH, Kurtz ZD, Cox LM, et al. Helminth Colonization Is Associated with Increased Diversity of the Gut Microbiota. PLoS Negl Trop Dis. 2014;8(5):e2880.

14. Cooper P, Walker AW, Reyes J, Chico M, Salter SJ, Vaca M, et al. Patent Human Infections with the Whipworm, Trichuris trichiura, Are Not Associated with Alterations in the Faecal Microbiota. PLoS One. 2013;8(10):e76573.

15. Cantacessi C, Giacomin P, Croese J, Zakrzewski M, Sotillo J, McCann L, et al. Impact of experimental hookworm infection on the human gut microbiota. J Infect Dis. 2014;210(9):1431–4.

16. Kay GL, Millard A, Sergeant MJ, Midzi N, Gwisai R, Mduluza T, et al. Differences in the Faecal Microbiome in Schistosoma haematobium Infected Children vs. Uninfected Children. PLoS Negl Trop Dis. 2015;9(6):e0003861.

17. Martin I, Djuardi Y, Sartono E, Rosa BA, Supali T, Mitreva M, et al. Dynamic changes in human-gut microbiome in relation to a placebo-controlled anthelminthic trial in Indonesia. PLoS Negl Trop Dis. 2018;12(8):e0006620.

18. Schneeberger PHH, Coulibaly JT, Gueuning M, Moser W, Coburn B, Frey JE, et al. Off-target effects of tribendimidine, tribendimidine plus ivermectin, tribendimidine plus oxantel-pamoate, and albendazole plus oxantel-pamoate on the human gut microbiota. Int J Parasitol Drugs Drug Resist. 2018;8(3):372–378.

19. Jovel J, Patterson J, Wang W, Hotte N, O’Keefe S, Mitchel T, et al. Characterization of the Gut Microbiome Using 16S or Shotgun Metagenomics. Front Microbiol. 2016;7:459.

20. Ranjan R, Rani A, Metwally A, McGee HS, Perkins DL. Analysis of the microbiome: Advantages of whole genome shotgun versus 16S amplicon sequencing. Biochem Biophys Res Commun. 2016;469(4):967–77.

21. Bowers RM, Kyrpides NC, Stepanauskas R, Harmon-Smith M, Doud D, Reddy TBK, et al. Minimum information about a single amplified genome (MISAG) and a metagenome-assembled genome (MIMAG) of bacteria and archaea. Nat Biotechnol. 2017;35(8):725–731.

22. Nayfach S, Shi ZJ, Seshadri R, Pollard KS, Kyrpides NC. New insights from uncultivated genomes of the global human gut microbiome. Nature. 2019;568(7753):505–510.

23. Pasolli E, Asnicar F, Manara S, Zolfo M, Karcher N, Armanini F, et al. Extensive Unexplored Human Microbiome Diversity Revealed by Over 150,000 Genomes from Metagenomes Spanning Age, Geography, and Lifestyle. Cell. 2019;176(3):649–662.e20.

24. Almeida A, Mitchell AL, Boland M, Forster SC, Gloor GB, Tarkowska A, et al. A new genomic blueprint of the human gut microbiota. Nature. 2019;568(7753):499–504.

25. Forster SC, Kumar N, Anonye BO, Almeida A, Viciani E, Stares MD, et al. A human gut bacterial genome and culture collection for improved metagenomic analyses. Nat Biotechnol. 2019;37(2):186–192.

26. Zou Y, Xue W, Luo G, Deng Z, Qin P, Guo R, et al. 1,520 reference genomes from cultivated human gut bacteria enable functional microbiome analyses. Nat Biotechnol. 2019;37(2):179–185.

27. Kim CY, Lee M, Yang S, Kim K, Yong D, Kim HR, et al. Human reference gut microbiome catalog including newly assembled genomes from under-represented Asian metagenomes. Genome Med. 2021;13(1):134.

28. Mallick H, Rahnavard A, McIver LJ, Ma S, Zhang Y, Nguyen LH, et al. Multivariable Association Discovery in Population-scale Meta-omics Studies. PLoS Comput Biol. 2021;17(11):e1009442.

29. Ajibola O, Rowan AD, Ogedengbe CO, Mshelia MB, Cabral DJ, Eze AA, et al. Urogenital schistosomiasis is associated with signatures of microbiome dysbiosis in Nigerian adolescents. Sci Rep. 2019;9(1):829.

30. Rubel MA, Abbas A, Taylor LJ, Connell A, Tanes C, Bittinger K, et al. Lifestyle and the presence of helminths is associated with gut microbiome composition in Cameroonians. Genome Biol. 2020;21(1):122.

31. Durazzi F, Sala C, Castellani G, Manfreda G, Remondini D, De Cesare A. Comparison between 16S rRNA and shotgun sequencing data for the taxonomic characterization of the gut microbiota. Sci Rep. 2021;11(1):3030.

32. Toro-Londono MA, Bedoya-Urrego K, Garcia-Montoya GM, Galvan-Diaz AL, Alzate JF. Intestinal parasitic infection alters bacterial gut microbiota in children. PeerJ. 2019;7:e6200.

33. Huwe T, Prusty BK, Ray A, Lee S, Ravindran B, Michael E. Interactions between Parasitic Infections and the Human Gut Microbiome in Odisha, India. Am J Trop Med Hyg. 2019;100(6):1486–1489.

34. Lee SC, Tang MS, Easton AV, Devlin JC, Chua LL, Cho I, et al. Linking the effects of helminth infection, diet and the gut microbiota with human whole-blood signatures. PLoS Pathog. 2019;15(12):e1008066.

35. Fierer N, Lennon JT. The generation and maintenance of diversity in microbial communities. Am J Bot. 2011;98(3):439–48.

36. Zhou D, Zhang H, Bai Z, Zhang A, Bai F, Luo X, et al. Exposure to soil, house dust and decaying plants increases gut microbial diversity and decreases serum immunoglobulin E levels in BALB/c mice. Environ Microbiol. 2016;18(5):1326–37.

37. Shyam Prasad G, Girisham S, Reddy SM. Microbial transformation of albendazole. Indian J Exp Biol. 2010;48(4):415–20.

38. Lanusse CE, Nare B, Gascon LH, Prichard RK. Metabolism of albendazole and albendazole sulphoxide by ruminal and intestinal fluids of sheep and cattle. Xenobiotica. 1992;22(4):419–26.

39. Moser W, Schindler C, Keiser J. Efficacy of recommended drugs against soil transmitted helminths: systematic review and network meta-analysis. BMJ. 2017;358:j4307.

40. Adisakwattana P, Yoonuan T, Phuphisut O, Poodeepiyasawat A, Homsuwan N, Gordon CA, et al. Clinical helminthiases in Thailand border regions show elevated prevalence levels using qPCR diagnostics combined with traditional microscopic methods. Parasit Vectors. 2020;13(1):416.

41. Montresor A, Crompton DWT, Hall A, Bundy DAP, Savioli L. Guidelines for the evaluation of soil-transmitted helminthiasis and schistosomiasis at community level. World Health Organization: Geneva. 1998.

42. Chin YT, Lim YA, Chong CW, Teh CS, Yap IK, Lee SC, et al. Prevalence and risk factors of intestinal parasitism among two indigenous sub-ethnic groups in Peninsular Malaysia. Infect Dis Poverty. 2016;5(1):77.

43. Langmead B, Salzberg SL. Fast gapped-read alignment with Bowtie 2. Nat Methods.2012;9(4):357–9.

44. Andrews S. FastQC: a quality control tool for high throughput sequence data. 2010.

45. Wood DE, Lu J, Langmead B. Improved metagenomic analysis with Kraken 2. Genome Biol. 2019;20(1):257.

46. Lu J, Breitwieser FP, Thielen P, Salzberg SL. Bracken: Estimating species abundance in metagenomics data. PeerJ Comput Sci. 2017;3:e104.

47. Wickham, H. ggplot2. tidyverse 2021. Available from: https://ggplot2.tidyverse.org/.

48. Kassambara A. rstatix: Pipe-Friendly Framework for Basic Statistical Tests. r-project 2021. https://cran.r-project.org/web/packages/rstatix/index.html. Accessed 11 October 2021.

49. McMurdie PJ, Holmes S. phyloseq: An R Package for Reproducible Interactive Analysis and Graphics of Microbiome Census Data. PLoS One. 2013;8(4):e61217.

50. Kassambara. A. ggpubr: ‘ggplot2’ Based Publication Ready Plots. datanovia 2021. https://rpkgs.datanovia.com/ggpubr/. Accessed 11 October 2021.

51. Brown CT, Izber L. sourmash: a library for MinHash sketching of DNA. J. Open Source Softw. 2016;1(5):27.

52. Breiman L, Cutler A, Liaw A, Breiman MW. randomForest: Breiman and Cutler’s Random Forests for Classification and Regression. r-project 2018. https://cran.r-project.org/web/packages/randomForest/index.html. Accessed on 11 October 2021.

53. Amunategui M. SMOTE - Supersampling Rare Events in R. Github 2014. http://amunategui.github.io/smote/#. Accessed on 18 October 2021.

54. Putri VM, Masjkur M, Suhaeni C. Performance of SMOTE in a random forest and naive Bayes classifier for imbalanced Hepatitis-B vaccination status. J Phys Conf Ser. 2021:1863(012073):9.

55. Brownlee J. Tune Machine Learning Algorithms in R (random forest case study). Machine Learning Mastery 2016. https://machinelearningmastery.com/tune-machine-learning-algorithms-in-r/. Accessed on 11 October 2021.

56. Douglas G. Random Forest Tutorial. LangilleLab/microbiome_helper. 2020. https://github.com/LangilleLab/microbiome_helper/wiki/Random-Forest-Tutorial. Accessed on 11 October 2021.

57. Humboldt AV, Bonpland A. Essay on the Geography of Plants. University of Chicago Press. 2010.

58. Shannon CE. A Mathematical Theory of Communication. Bell Syst Tech J. 1948;27:379–423, 623–656.

59. Simpson EH. Measurement of Diversity. Nature. 1949;163.

60. Ssekagiri A, Sloan WT, Ijaz UZ. microbiomeSeq: An R package for microbial community analysis in an environmental context. University of Glasgow : School of Engineering 2017. http://userweb.eng.gla.ac.uk/umer.ijaz/projects/microbiomeSeq_Tutorial.html. Accessed on 18 October 2021.

61. Wilcoxon F. Individual Comparisons by Ranking Methods. Biometrics. 1945;1(6):80–83.

62. Spearman C. Demonstration of Formulæ for True Measurement of Correlation. Am J Psychol. 1907;18(2):161–169.

63. Boogaart KG, Tolosana-Delgado R, Bren M. compositions: Compositional Data Analysis. r-project 2021. https://cran.r-project.org/web/packages/compositions/index.html. Accessed on 11 October 2021.

64. Anderson MJ. Distance-based tests for homogeneity of multivariate dispersions. Biometrics. 2006;62(1):245–253.

65. Clarke KR. Non-parametric multivariate analyses of changes in community structure. Austral Ecol. 1993;18(1):117–143.

66. Oksanen J, Blanchet FG, Friendly M, Kindt R, Legendre P, McGlinn D, et al. vegan: Community Ecology Package. r-project 2020. https://cran.r-project.org/web/packages/vegan/index.html.

67. Lin H, Peddada SD. Analysis of compositions of microbiomes with bias correction. Nat Commun. 2020;11(1):3514.

68. Emiola A, Oh J. High throughput in situ metagenomic measurement of bacterial replication at ultra-low sequencing coverage. Nat Commun. 2018;9(1):4956.

69. Bolger AM, Lohse M, Usadel B. Trimmomatic: a flexible trimmer for Illumina sequence data. Bioinformatics. 2014;30(15):2114–20.

70. Hong C, Manimaran S, Shen Y, Perez-Rogers JF, Byrd AL, Castro-Nallar E, et al. PathoScope 2.0: a complete computational framework for strain identification in environmental or clinical sequencing samples. Microbiome. 2014;2:33.

