## Supplementary_Materials for "Gut Microbiome of Helminth Infected Indigenous Malaysians Is Context Dependent"

Supplementary Figure S1 to S21

Supplementary Table S1 to S4

**Fig. S1**

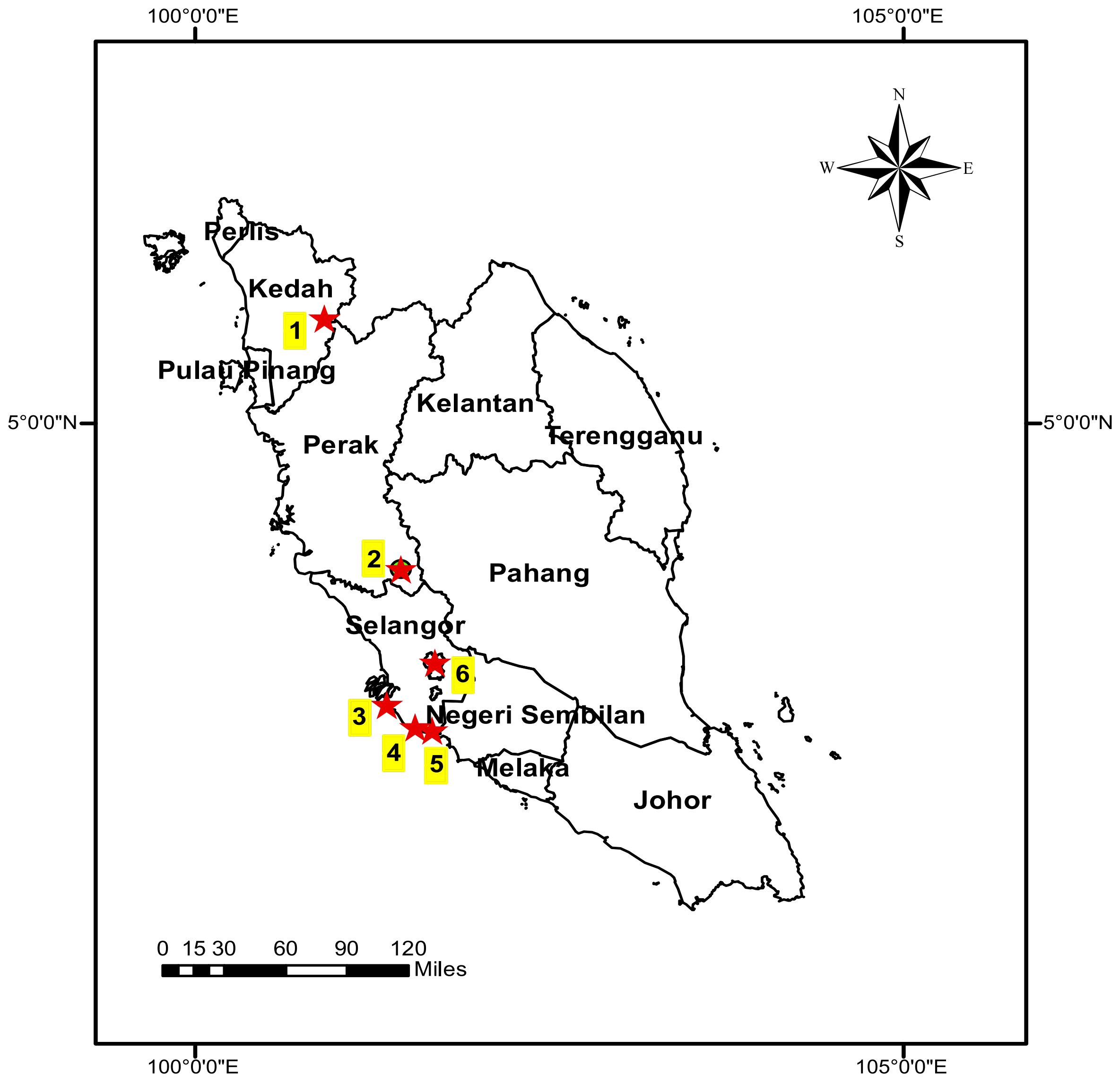


| No. | Villages | States | Tribes | Subtribes |
|---|---|---|---|---|
| 1 | Legong | Kedah | Semang/ Negrito | Kensiu |
| 2 | Rasau | Perak | Senoi | Semai |
| 3 | Judah | Selangor | Senoi | Mah Meri |
| 4 | Sepat | Selangor | Senoi | Mah Meri |
| 5 | Bangkong | Selangor | Senoi | Mah Meri |
| 6 | KL | Kuala Lumpur | NA | NA |

**Fig. S1**

A geographic map showing the locations of each village and the Kuala Lumpur city in Peninsular Malaysia (stars and numbers) together with a table with other information including states, tribes and subtribes.

**Fig. S2**

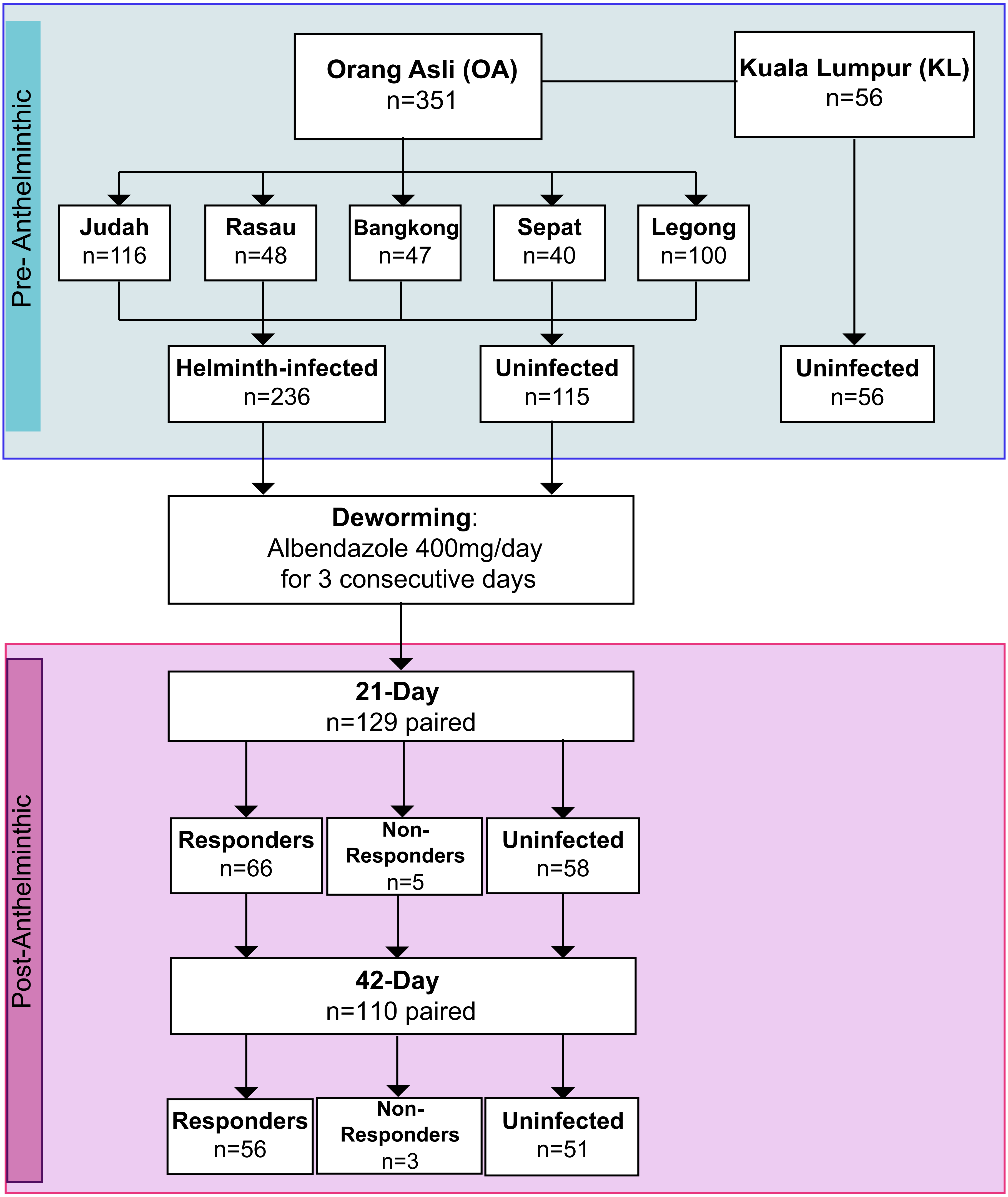
Pre- Anthelminthic
Orang Asli (OA)
n=351
Kuala Lumpur (KL)
n=56
Judah
n=116
Rasau
n=48
Bangkong
n=47
Sepat
n=40
Legong
n=100
Helminth-infected
n=236
Uninfected
n=115
Uninfected
n=56
Deworming:
Albendazole 400mg/day
for 3 consecutive days
Post-Anthelminthic
21-Day
n=129 paired
Responders
n=66
Non-
Responders
n=5
Uninfected
n=58
42-Day
n=110 paired
Responders
n=56
Non-
Responders
n=3
Uninfected
n=51

**Fig. S2**

A flow diagram of the total subjects (Orang Asli and urban citizens from Kuala Lumpur) involved in both the Pre-anthelminthic and Post-anthelminthic of this study.

### Fig. S3

13,273,611,903 paired end reads
(mean: 20,420,941±3,294,098;
max: 35,530,200; min: 13,037,828)

Raw reads

**1**

KneadData

11,480,206,516 paired end reads
(mean: 17,661,856 ±2,977,678.7;
max: 31,327,456; min: 6,683,658)

QC remove human read
adapter

**2**

Kracken 2
Bracken 2

Map to RefSeq, UHGG ,
HRGM database for
taxonomic classification

Downstream
analysis

**A**

Bray-Curtis and Jaccard:
PCoA, Adonis, ANOSIM
Beta-dispersion

**B**

Alpha diversity:
Species Richness,
Shannon, Simpson

**C**

Differential abundance:
MaAsLin2, ANCOM

**3**

Sourmash

K-mer based approach

**4**

GRiD

Estimation of the growth
rate of bacteria

**Created with BioRender.com**

**Fig. S3**

A flow diagram summarizing the bioinformatic analysis from raw reads, 1) Quality filtering, remove human reads and adapter (KneadData), taxonomic classification (Kraken2 & Bracken2), 3) K-mer based approach (Sourmash) and 4) Estimation of bacterial growth rate (GRiD) to downstream analysis (A-C) such as beta-diversity, alpha diversity and differential abundance.

**Fig. S4**

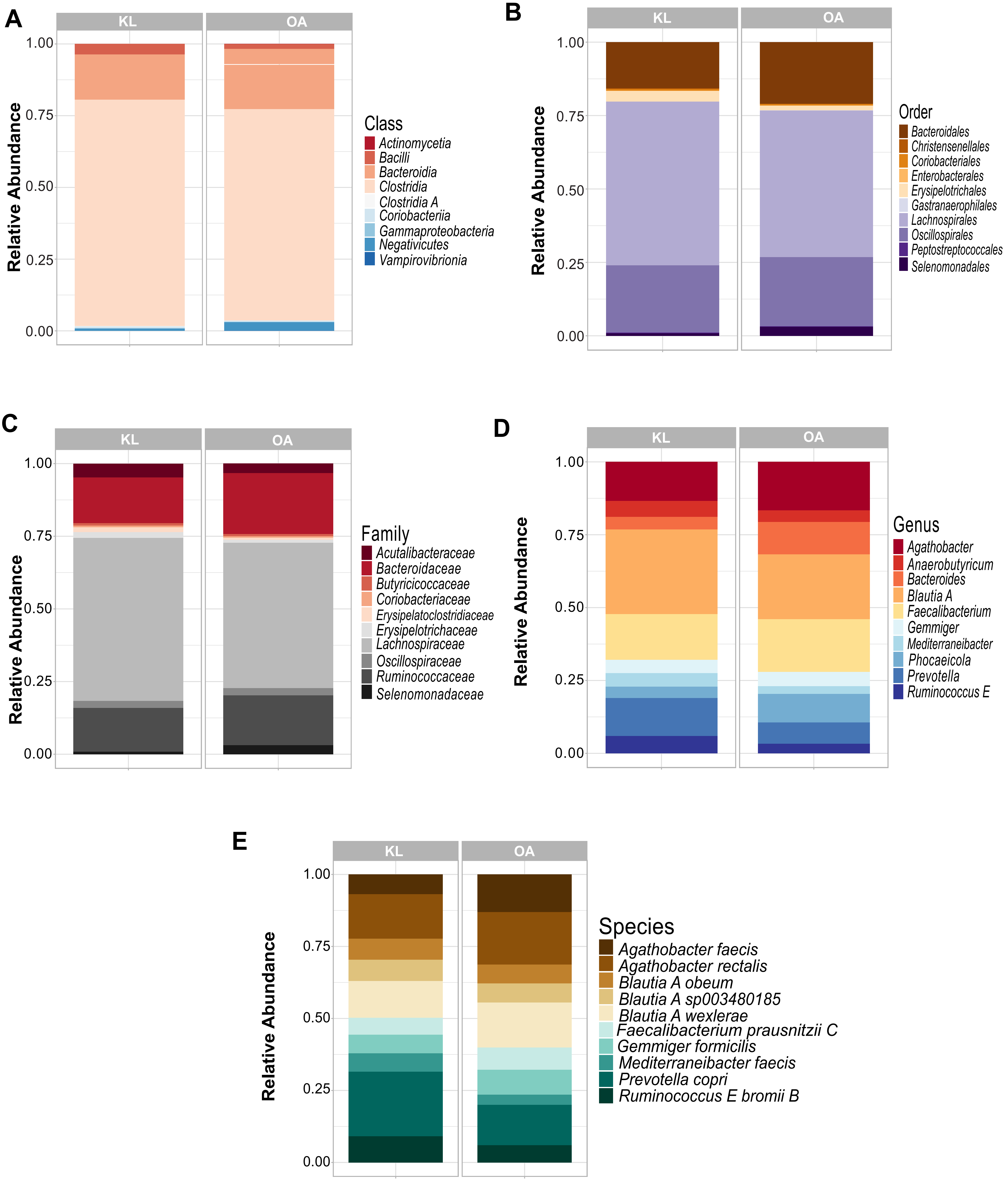
A
KL
OA
Relative Abundance
1.00
0.75
0.50
0.25
0.00
Class
Actinomycetia
Bacilli
Bacteroidia
Clostridia
Clostridia A
Coriobacteriia
Gammaproteobacteria
Negativicutes
Vampirovibrionia
B
KL
OA
Relative Abundance
1.00
0.75
0.50
0.25
0.00
Order
Bacteroidales
Christensenellales
Coriobacteriales
Enterobacterales
Erysipelotrichales
Gastranaerophilales
Lachnospirales
Oscillospirales
Peptostreptococcales
Selenomonadales
C
KL
OA
Relative Abundance
1.00
0.75
0.50
0.25
0.00
Family
Acutalibacteraceae
Bacteroidaceae
Butyricicoccaceae
Coriobacteriaceae
Erysipelatoclostridiaceae
Erysipelotrichaceae
Lachnospiraceae
Oscillospiraceae
Ruminococcaceae
Selenomonadaceae
D
KL
OA
Relative Abundance
1.00
0.75
0.50
0.25
0.00
Genus
Agathobacter
Anaerobutyricum
Bacteroides
Blautia A
Faecalibacterium
Gemmiger
Mediterraneibacter
Phocaeicola
Prevotella
Ruminococcus E
E
KL
OA
Relative Abundance
1.00
0.75
0.50
0.25
0.00
Species
Agathobacter faecis
Agathobacter rectalis
Blautia A obeum
Blautia A sp003480185
Blautia A wexlerae
Faecalibacterium prausnitzii C
Gemmiger formicilis
Mediterraneibacter faecis
Prevotella copri
Ruminococcus E bromii B

**Fig. S4**

Difference in the composition of core microbiota between Orang Asli cohort and KL cohort in different taxonomic rank, which include: **A** Class, **B** Order, **C** Family, **D** Genus & **E** Species.

Fig. S5

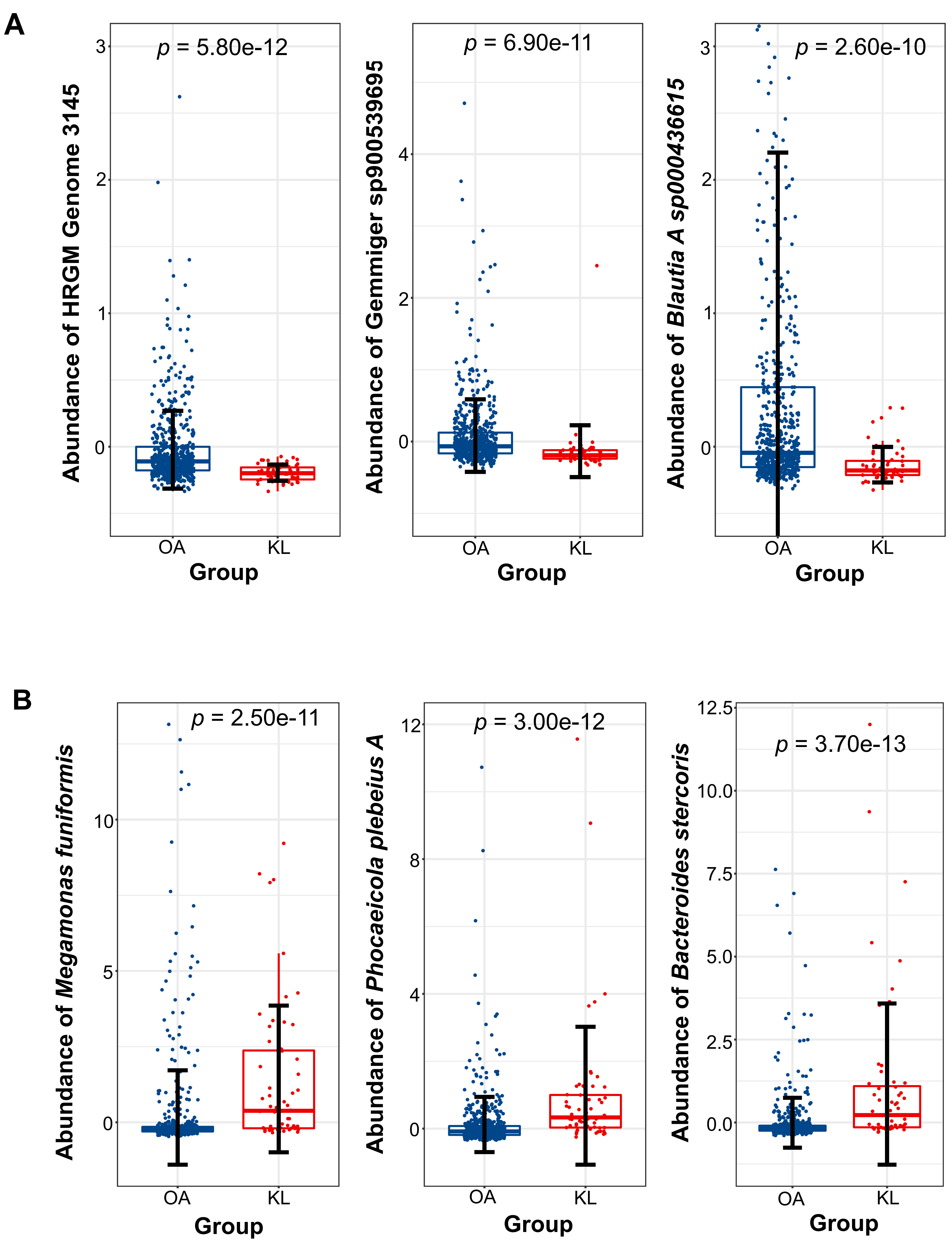
A
p = 5.80e-12
Abundance of HRGM Genome 3145
3
2
1
0
OA
KL
Group
p = 6.90e-11
Abundance of Gemmiger sp900539695
4
2
0
OA
KL
Group
p = 2.60e-10
Abundance of Blautia A sp000436615
3
2
1
0
OA
KL
Group
B
p = 2.50e-11
Abundance of Megamonas funiformis
10
5
0
OA
KL
Group
p = 3.00e-12
Abundance of Phocaeicola plebeius A
12
8
4
0
OA
KL
Group
p = 3.70e-13
Abundance of Bacteroides stercoris
12.5
10.0
7.5
5.0
2.5
0.0
OA
KL
Group

**Fig. S5**

Box plots displaying the selected core microbial species that have high variation between Orang Asli (OA) cohort and Kuala Lumpur (KL) cohort based on the random forest analysis. The relative abundances of core microbial species between Orang Asli cohort and KL cohort was tested using Wilcoxon rank sum test. **A** Species with significant higher abundance in Orang Asli cohort than KL cohort, which include (from left to right): HRGM Genome 3145, *Gemmiger sp900539695* and *Blautia A sp00043661,* respectively. **B** Species with significant higher abundance in KL cohort than the Orang Asli cohort, which include (from left to right): *Megamonas funiformis*, *Phocaeicola plebeius A* and *Bacteroides stercoris,* respectively.

Fig. S6

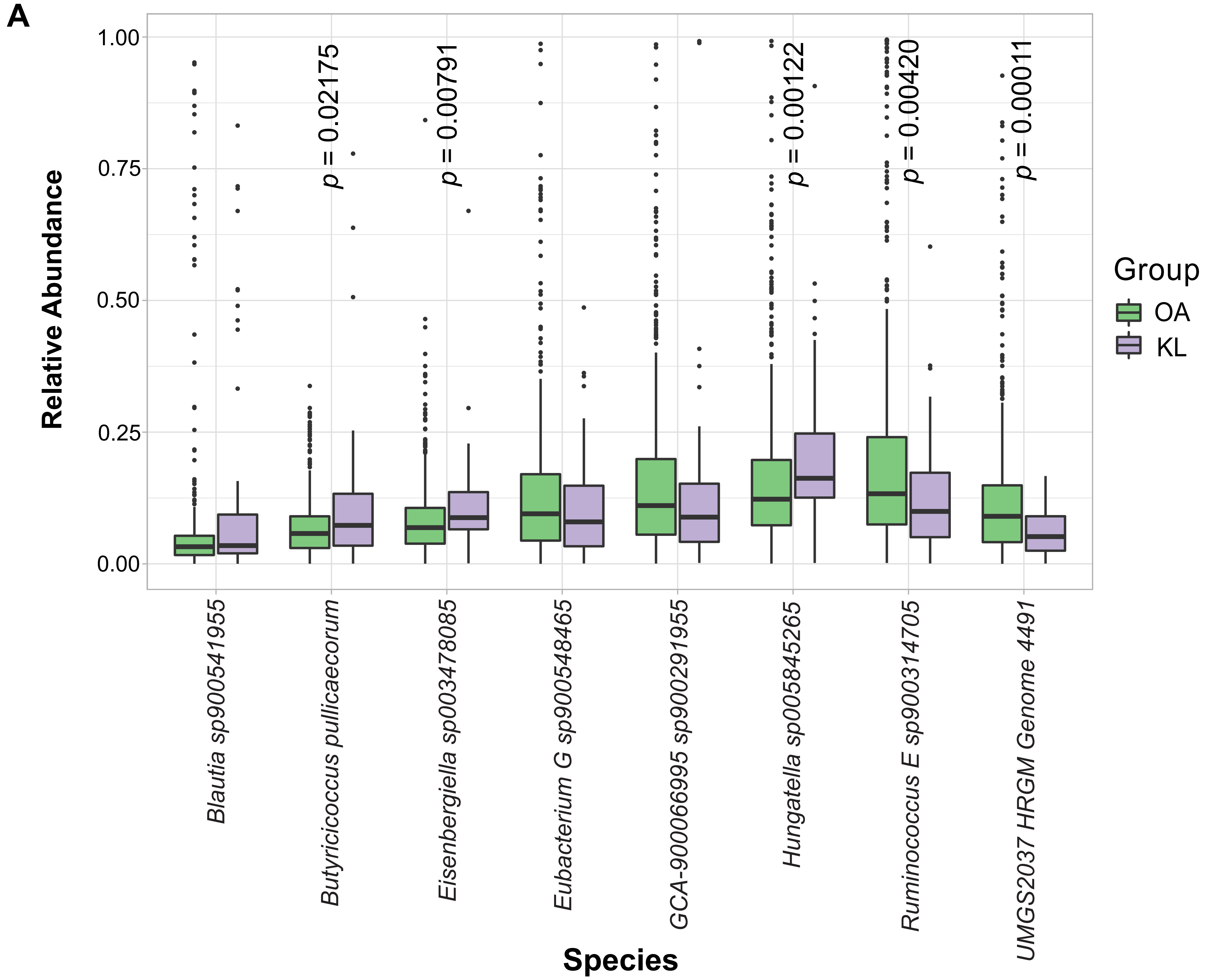

A
Relative Abundance
1.00
0.75
0.50
0.25
0.00
p = 0.02175
p = 0.00791
p = 0.00122
p = 0.00420
p = 0.00011
Group
OA
KL
Blautia sp900541955
Butyricicoccus pullicaecorum
Eisenbergiella sp003478085
Eubacterium G sp900548465
GCA-900066995 sp900291955
Hungatella sp005845265
Ruminococcus E sp900314705
UMGS2037 HRGM Genome 4491
Species


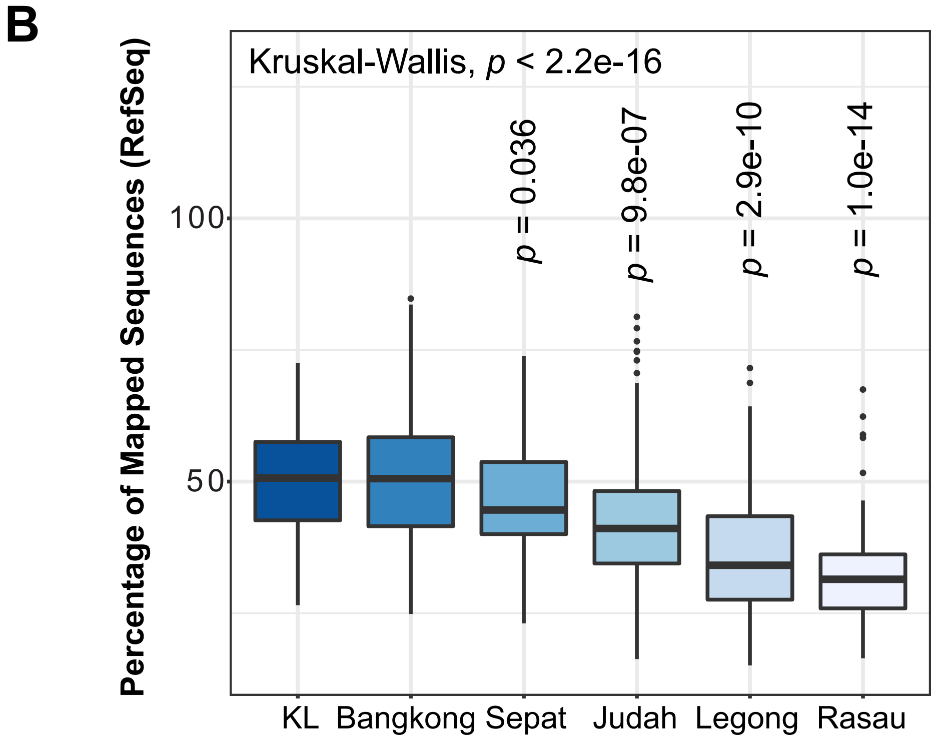

B
Percentage of Mapped Sequences (RefSeq)
Kruskal-Wallis, p < 2.2e-16
100
50
p = 0.036
p = 9.8e-07
p = 2.9e-10
p = 1.0e-14
KL
Bangkong
Sepat
Judah
Legong
Rasau


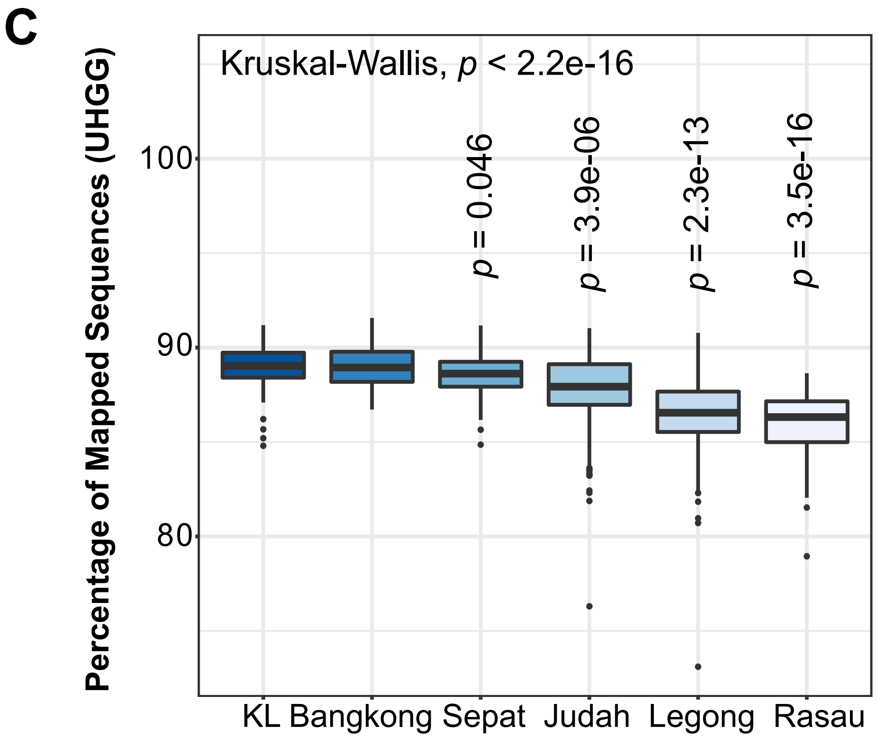

C
Percentage of Mapped Sequences (UHGG)
Kruskal-Wallis, p < 2.2e-16
100
90
80
p = 0.046
p = 3.9e-06
p = 2.3e-13
p = 3.5e-16
KL
Bangkong
Sepat
Judah
Legong
Rasau

**Fig. S6**

Effects of geographical location on core gut microbiota and the percentage of unmapped reads in the microbiome. **A** The core gut microbial species showing the largest variation (cut-off 6.0 for the coefficient of variation) between Orang Asli and Kuala Lumpur cohort in Malaysia across 650 samples. Box plots illustrate the percentage of mapped reads in **B** RefSeq (i.e., Bacteria, protozoa, fungi, viral, archaea) database, and **C** Unified Human Gastrointestinal Genome (UHGG) database in different geographical locations. Pairwise comparison between each village and the KL cohort was tested using Wilcoxon rank sum test whereas the comparison for all groups was tested using Kruskal-Wallis.

Fig. S7

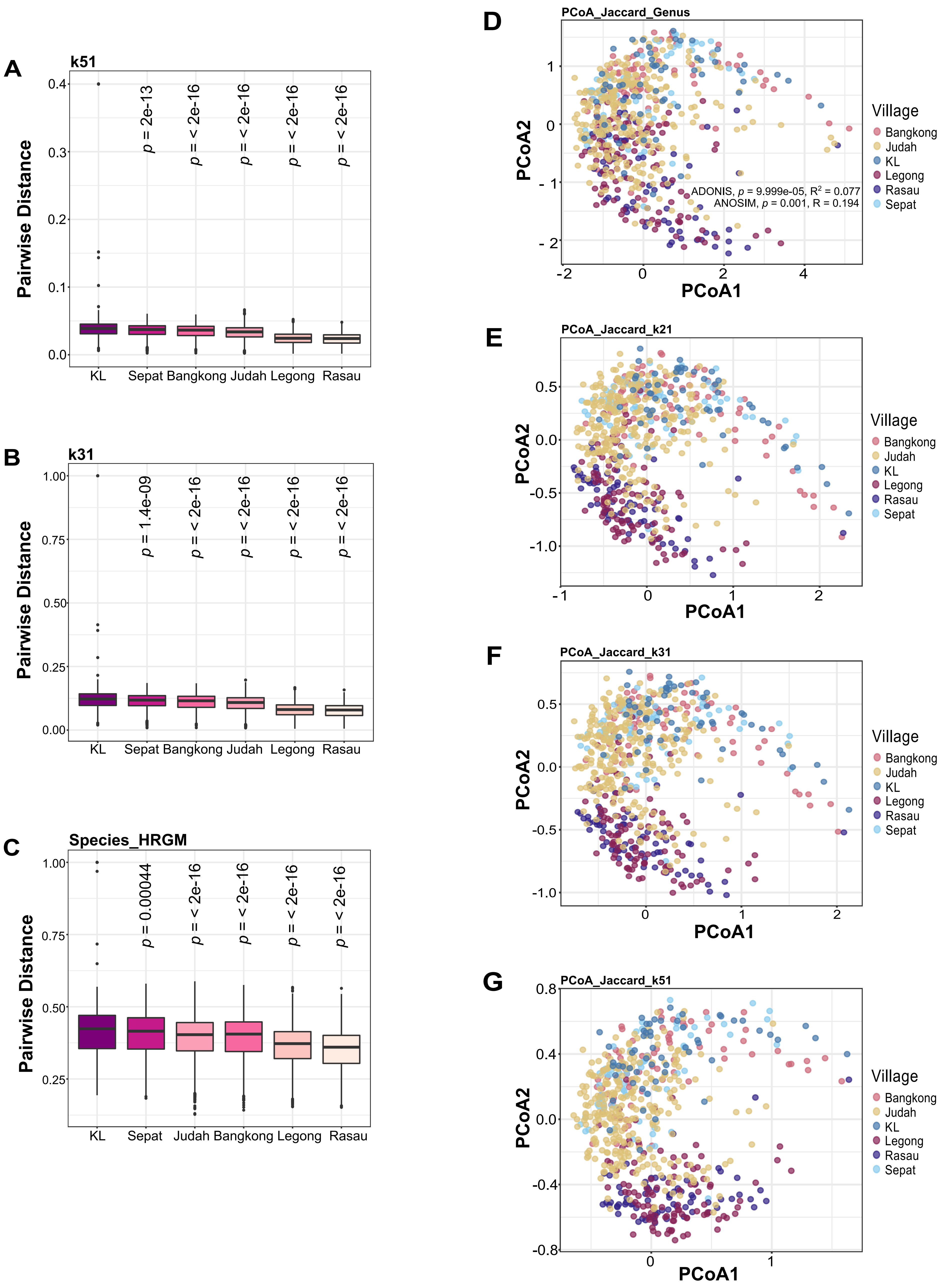
A
k51
Pairwise Distance
p = 2e-13
p = < 2e-16
p = < 2e-16
p = < 2e-16
p = < 2e-16
KL Sepat Bangkong Judah Legong Rasau
B
k31
Pairwise Distance
p = 1.4e-09
p = < 2e-16
p = < 2e-16
p = < 2e-16
p = < 2e-16
KL Sepat Bangkong Judah Legong Rasau
C
Species_HRGM
Pairwise Distance
p = 0.00044
p = < 2e-16
p = < 2e-16
p = < 2e-16
p = < 2e-16
KL Sepat Judah Bangkong Legong Rasau
D
PCoA_Jaccard_Genus
PCoA1
PCoA2
ADONIS, p = 9.999e-05, R2 = 0.077
ANOSIM, p = 0.001, R = 0.194
Village
Bangkong
Judah
KL
Legong
Rasau
Sepat
E
PCoA_Jaccard_k21
PCoA1
PCoA2
F
PCoA_Jaccard_k31
PCoA1
PCoA2
G
PCoA_Jaccard_k51
PCoA1
PCoA2

**Fig.S7**

Beta diversity of 650 samples [Orang Asli (OA) and Kuala Lumpur (KL) cohort]. Comparison of pairwise beta diversity of all villages to KL cohort, assessed by Jaccard distance based on distance of **A** nucleotide k-mer sketches (k=51), **B** k-mer sketches (k=31), and **C** species level. Pairwise comparison between each village against the KL cohort was tested using Wilcoxon rank sum test. Principal Coordinates Analysis (PCoA) of Jaccard distance based on **D** genus, **E** k-mer sketches =21, **F** k-mer sketches =31, and **G** k-mer sketches =51 in OA and KL cohort. The individuals from different geographical locations were denoted by different colors.

### Fig. S8

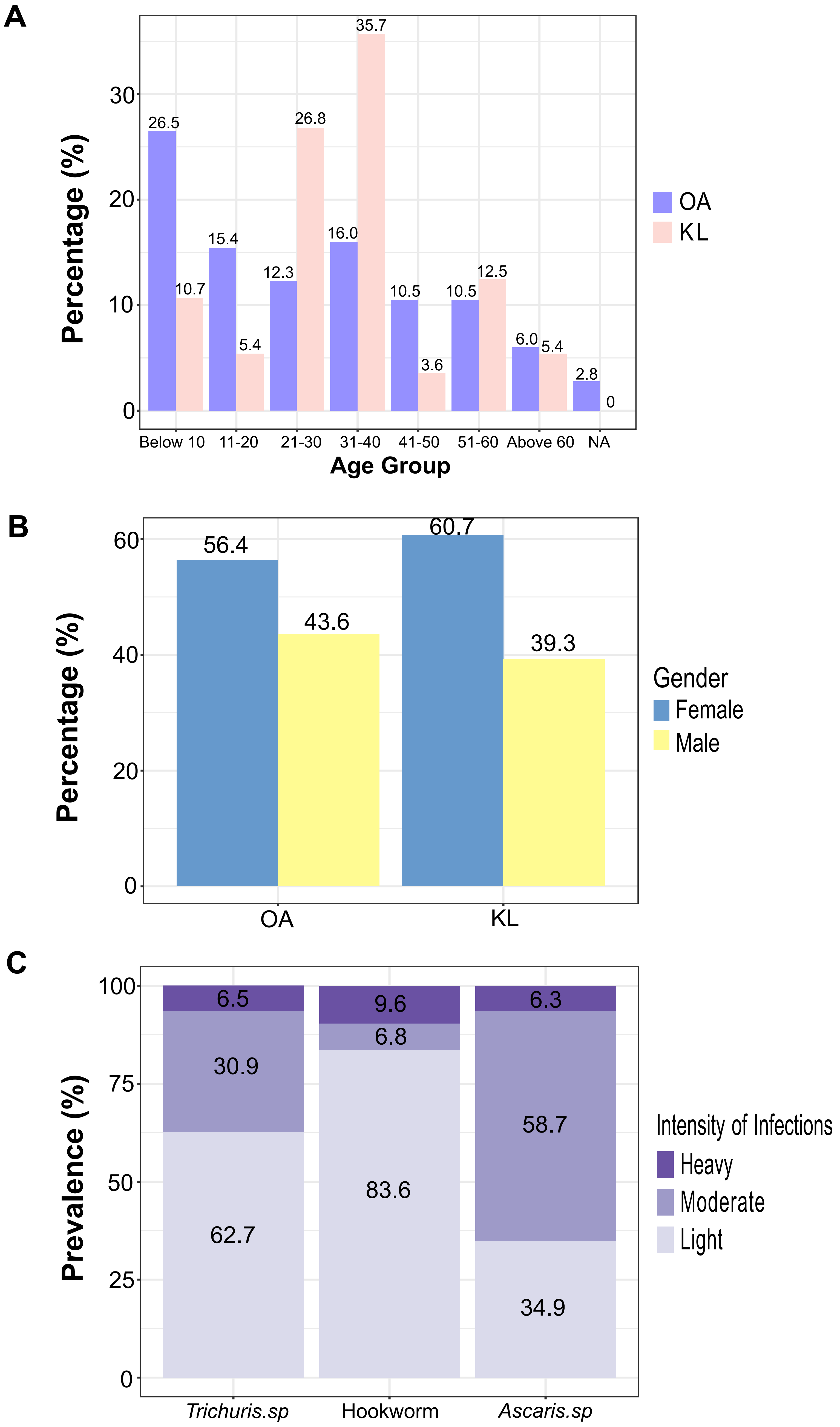
A
Percentage (%)
0
10
20
30
26.5
10.7
15.4
5.4
12.3
26.8
16.0
35.7
10.5
3.6
10.5
12.5
6.0
5.4
2.8
0
Below 10
11-20
21-30
31-40
41-50
51-60
Above 60
NA
Age Group
OA
KL
B
Percentage (%)
0
20
40
60
56.4
43.6
60.7
39.3
OA
KL
Gender
Female
Male
C
Prevalence (%)
0
25
50
75
100
6.5
30.9
62.7
9.6
6.8
83.6
6.3
58.7
34.9
Trichuris.sp
Hookworm
Ascaris.sp
Intensity of Infections
Heavy
Moderate
Light

**Fig.S8**

Epidemiology data of the Orang Asli (OA) and Kuala Lumpur (KL) Cohort. **A** Distribution of the age group from OA and KL cohort, the OA and KL cohort were denoted by purple and pink color respectively. **B** Distribution of the gender from OA and KL cohort, the female and male cohort were denoted by blue and yellow color respectively**. D** The prevalence of different types of helminthiases, which include *Trichuris* infection, *Ascaris* infection and hookworm infection, the heavy, moderate and light infection were detonated by different purple color intensity.

**Fig. S9**

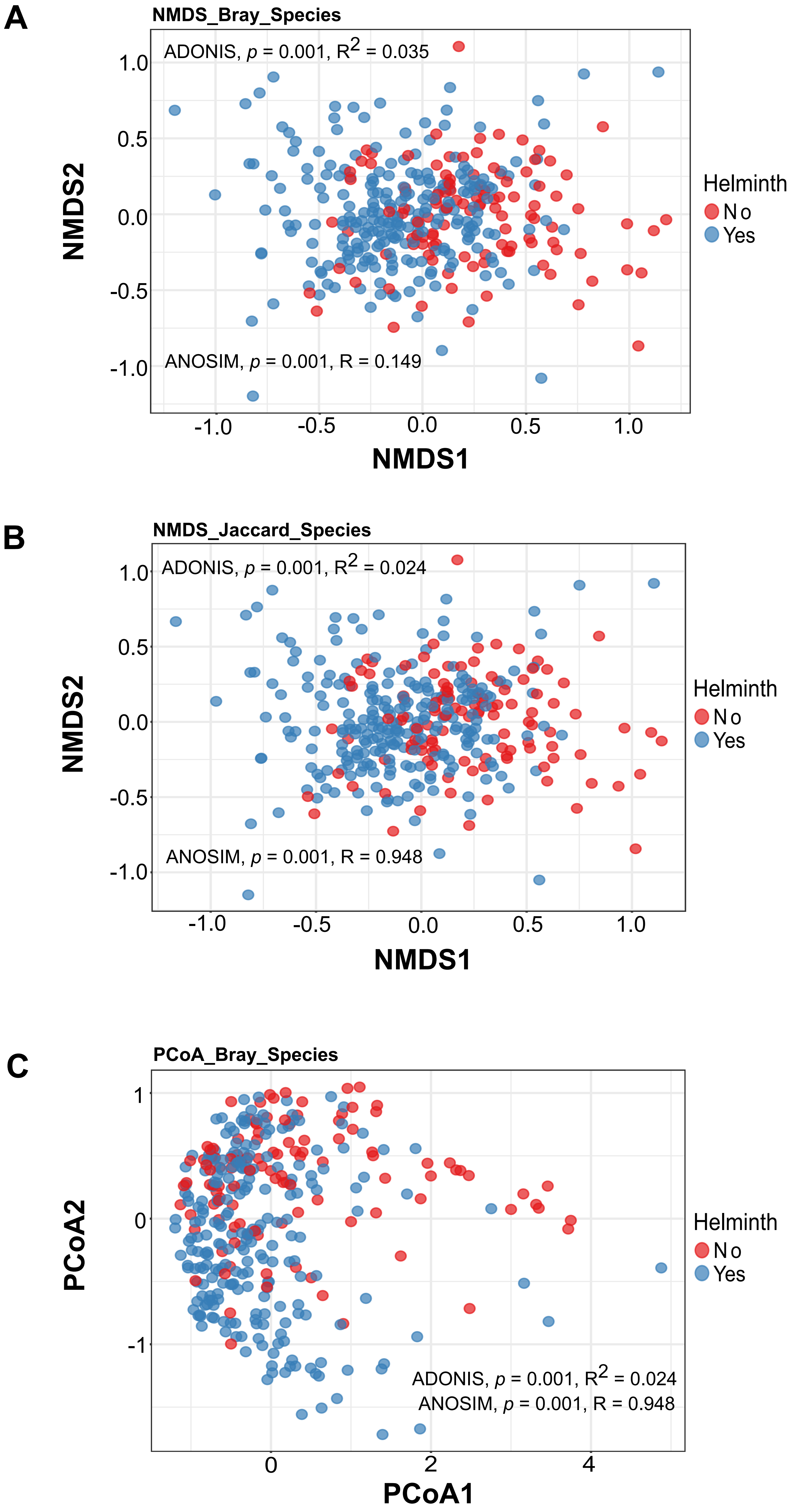

A
NMDS_Bray_Species
ADONIS, p = 0.001, R2 = 0.035
ANOSIM, p = 0.001, R = 0.149
NMDS1
NMDS2
Helminth
No
Yes
B
NMDS_Jaccard_Species
ADONIS, p = 0.001, R2 = 0.024
ANOSIM, p = 0.001, R = 0.948
NMDS1
NMDS2
Helminth
No
Yes
C
PCoA_Bray_Species
ADONIS, p = 0.001, R2 = 0.024
ANOSIM, p = 0.001, R = 0.948
PCoA1
PCoA2
Helminth
No
Yes

**Fig. S9**

Beta diversity comparing the gut microbiome between intestinal helminth infected and uninfected Orang Asli cohorts visualized using Non-metric multidimensional scaling (NMDS) plot of **A** Bray-curtis (ADONIS: p=0.001, $R^2$=0.035; ANOSIM: p=0.001, R=0.149) and **B** Jaccard distance (ADONIS: p=0.001, $R^2$=0.024; ANOSIM: p=0.001, R=0.948) and **C** Principal Coordinates Analysis (PCoA) of Jaccard distance (ADONIS: p=0.001, $R^2$=0.024; ANOSIM: p=0.001, R=0.948). The individuals infected and uninfected with intestinal helminths denoted by blue and red color respectively.

### Fig. S10

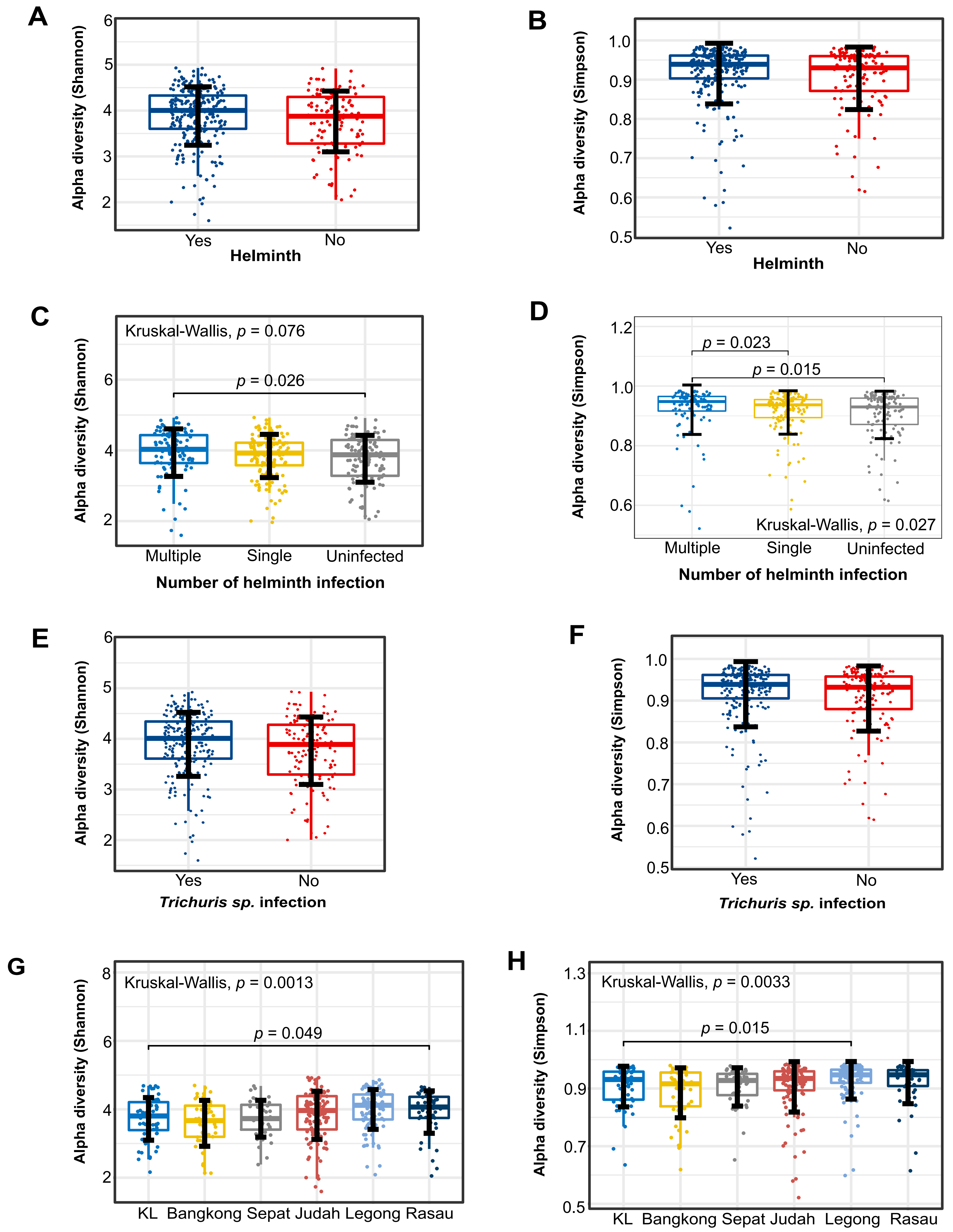
A
Alpha diversity (Shannon)
Yes
No
Helminth
B
Alpha diversity (Simpson)
Yes
No
Helminth
C
Kruskal-Wallis, p = 0.076
p = 0.026
Alpha diversity (Shannon)
Multiple
Single
Uninfected
Number of helminth infection
D
p = 0.023
p = 0.015
Kruskal-Wallis, p = 0.027
Alpha diversity (Simpson)
Multiple
Single
Uninfected
Number of helminth infection
E
Alpha diversity (Shannon)
Yes
No
Trichuris sp. infection
F
Alpha diversity (Simpson)
Yes
No
Trichuris sp. infection
G
Kruskal-Wallis, p = 0.0013
p = 0.049
Alpha diversity (Shannon)
KL
Bangkong
Sepat
Judah
Legong
Rasau
H
Kruskal-Wallis, p = 0.0033
p = 0.015
Alpha diversity (Simpson)
KL
Bangkong
Sepat
Judah
Legong
Rasau

**Fig. S10**

Box plots showing alpha diversity of gut microbiome profile using **A** Shannon diversity and **B** Simpson diversity index on individuals infected and uninfected with intestinal helminths; **C** Shannon diversity and **D** Simpson diversity index on different numbers of intestinal helminth infection; **E** Shannon diversity and **F** Simpson diversity index on individuals infected and uninfected with *Trichuris* sp. infection; **G** Shannon diversity and **H** Simpson diversity index on different villages. The statistical difference between two groups was tested using the Wilcoxon rank sum test whereas more than two groups was tested using Kruskal-Wallis.

### Fig. S11

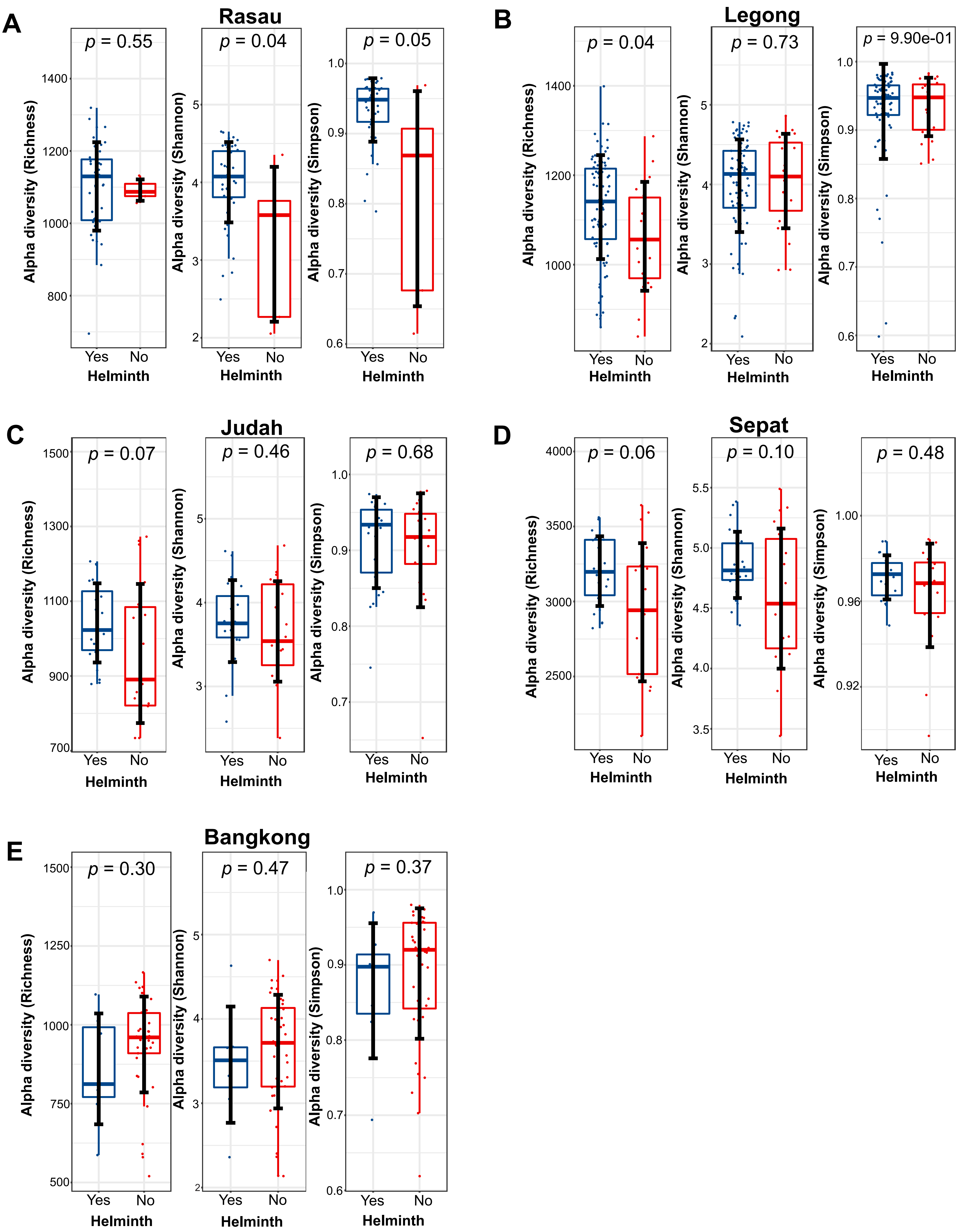
A
Rasau
p = 0.55
p = 0.04
p = 0.05
B
Legong
p = 0.04
p = 0.73
p = 9.90e-01
C
Judah
p = 0.07
p = 0.46
p = 0.68
D
Sepat
p = 0.06
p = 0.10
p = 0.48
E
Bangkong
p = 0.30
p = 0.47
p = 0.37
Alpha diversity (Richness)
Alpha diversity (Shannon)
Alpha diversity (Simpson)
Yes
No
Helminth

**Fig. S11**

Box plots showing alpha diversity (i.e., Richness, Shannon and Simpson diversity index) of gut microbiome profile on Orang Asli that infected and uninfected with intestinal helminths from **A** Rasau, **B** Legong, **C** Judah, **D** Sepat, and **E** Bangkong. The comparison of the alpha diversity index between helminth-infected and non-infected samples is tested using the Wilcoxon rank sum test.

### Fig. S12

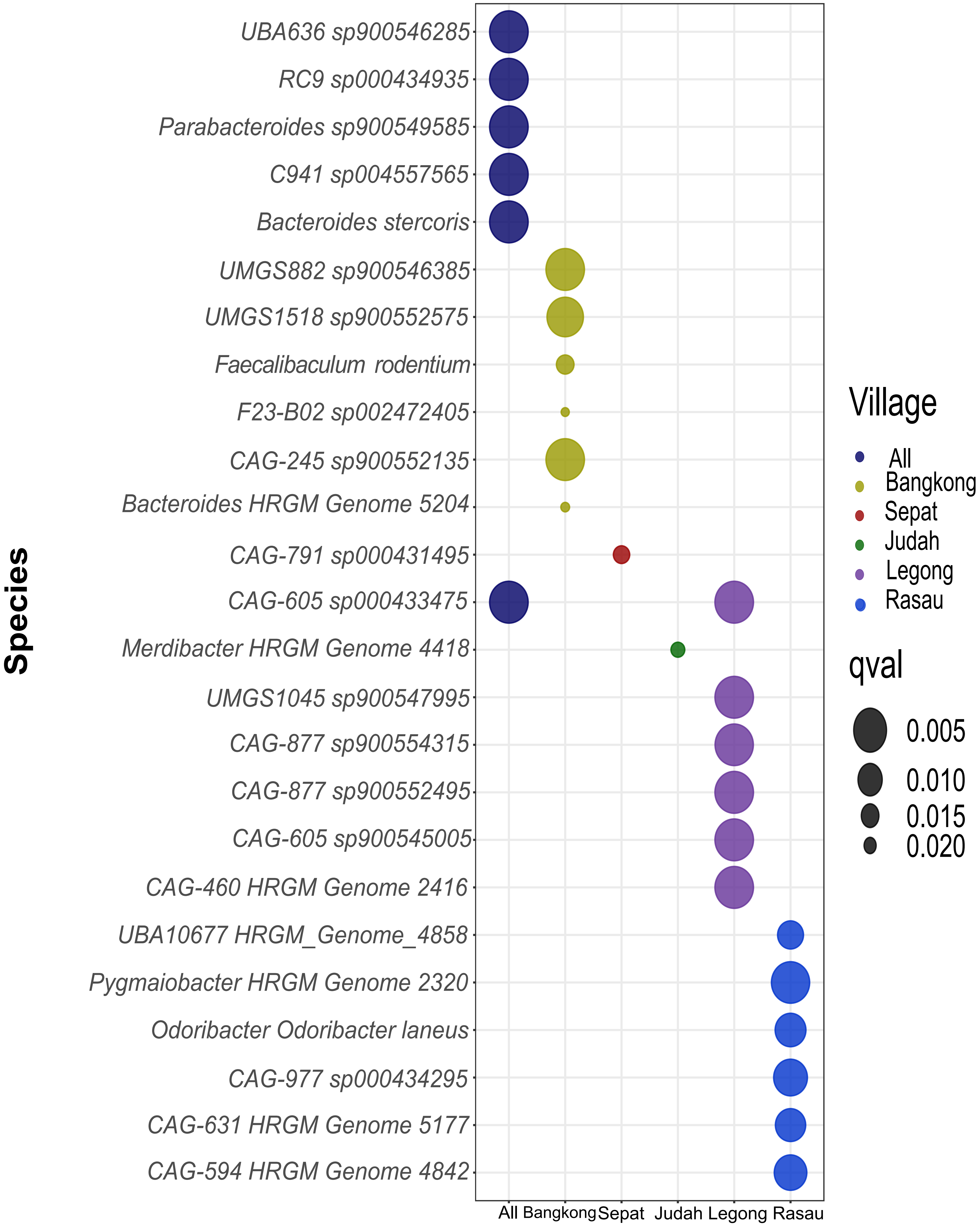

UBA636 sp900546285
RC9 sp000434935
Parabacteroides sp900549585
C941 sp004557565
Bacteroides stercoris
UMGS882 sp900546385
UMGS1518 sp900552575
Faecalibaculum rodentium
F23-B02 sp002472405
CAG-245 sp900552135
Bacteroides HRGM Genome 5204
CAG-791 sp000431495
CAG-605 sp000433475
Merdibacter HRGM Genome 4418
UMGS1045 sp900547995
CAG-877 sp900554315
CAG-877 sp900552495
CAG-605 sp900545005
CAG-460 HRGM Genome 2416
UBA10677 HRGM_Genome_4858
Pygmaiobacter HRGM Genome 2320
Odoribacter Odoribacter laneus
CAG-977 sp000434295
CAG-631 HRGM Genome 5177
CAG-594 HRGM Genome 4842
Species
All
Bangkong
Sepat
Judah
Legong
Rasau
Village
All
Bangkong
Sepat
Judah
Legong
Rasau
qval
0.005
0.010
0.015
0.020

**Fig. S12**

Bubble plot shows bacterial species that are differentially abundant between *Trichuris* infected and uninfected groups in all samples, as well as specific villages based on the output of the Analysis of Compositions of Microbiomes with Bias Correction (ANCOM-BC). The size of the bubble is negatively proportional to the p-value. The larger the bubble size displaying the lower p-value.

**Fig. S13**

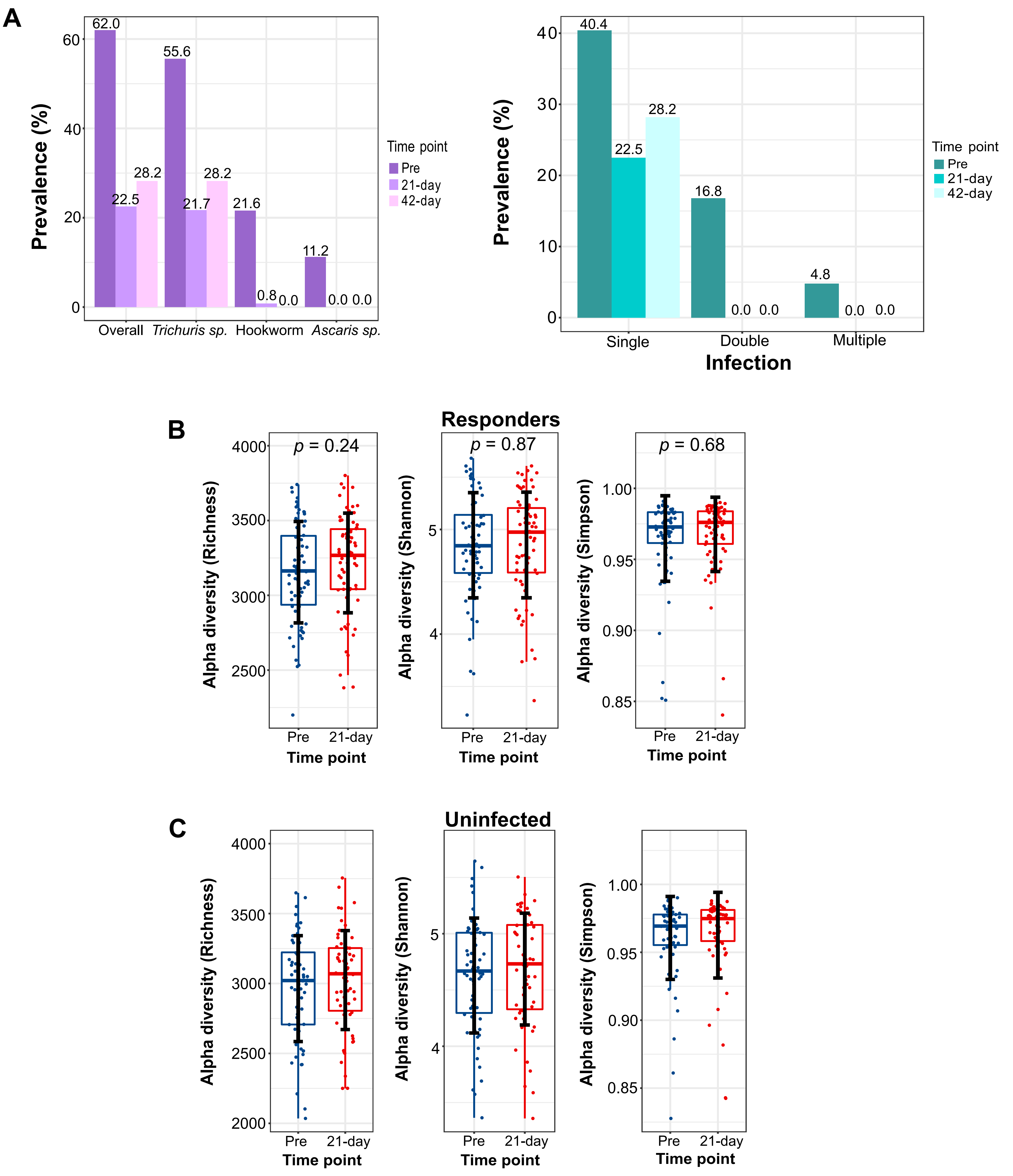
A
Prevalence (%)
62.0
22.5
28.2
55.6
21.7
28.2
21.6
0.8
0.0
11.2
0.0 0.0
Overall
Trichuris sp.
Hookworm
Ascaris sp.
Time point
Pre
21-day
42-day
40.4
22.5
28.2
16.8
0.0 0.0
4.8
0.0 0.0
Single
Double
Multiple
Infection
B
Responders
p = 0.24
p = 0.87
p = 0.68
Alpha diversity (Richness)
Alpha diversity (Shannon)
Alpha diversity (Simpson)
Pre
21-day
Time point
C
Uninfected
Alpha diversity (Richness)
Alpha diversity (Shannon)
Alpha diversity (Simpson)
Pre
21-day
Time point

**Fig. S13**

**A** Bar chart shows the changes in the prevalence of different types of helminthic infections in (Left), and the prevalence of the number of helminthic infections (Right) among the Pre-anthelminthic, 21-Day and 42-Day Post-anthelminthic. Boxplot showing alpha diversity (i.e., Richness, Shannon and Simpson diversity index) of gut microbiome profile on the **B** Responder, and **C** Uninfected. The comparison of the alpha diversity index between helminth-infected and non-infected samples is tested using the Wilcoxon sign rank test.

### Fig. S14

#### Responders

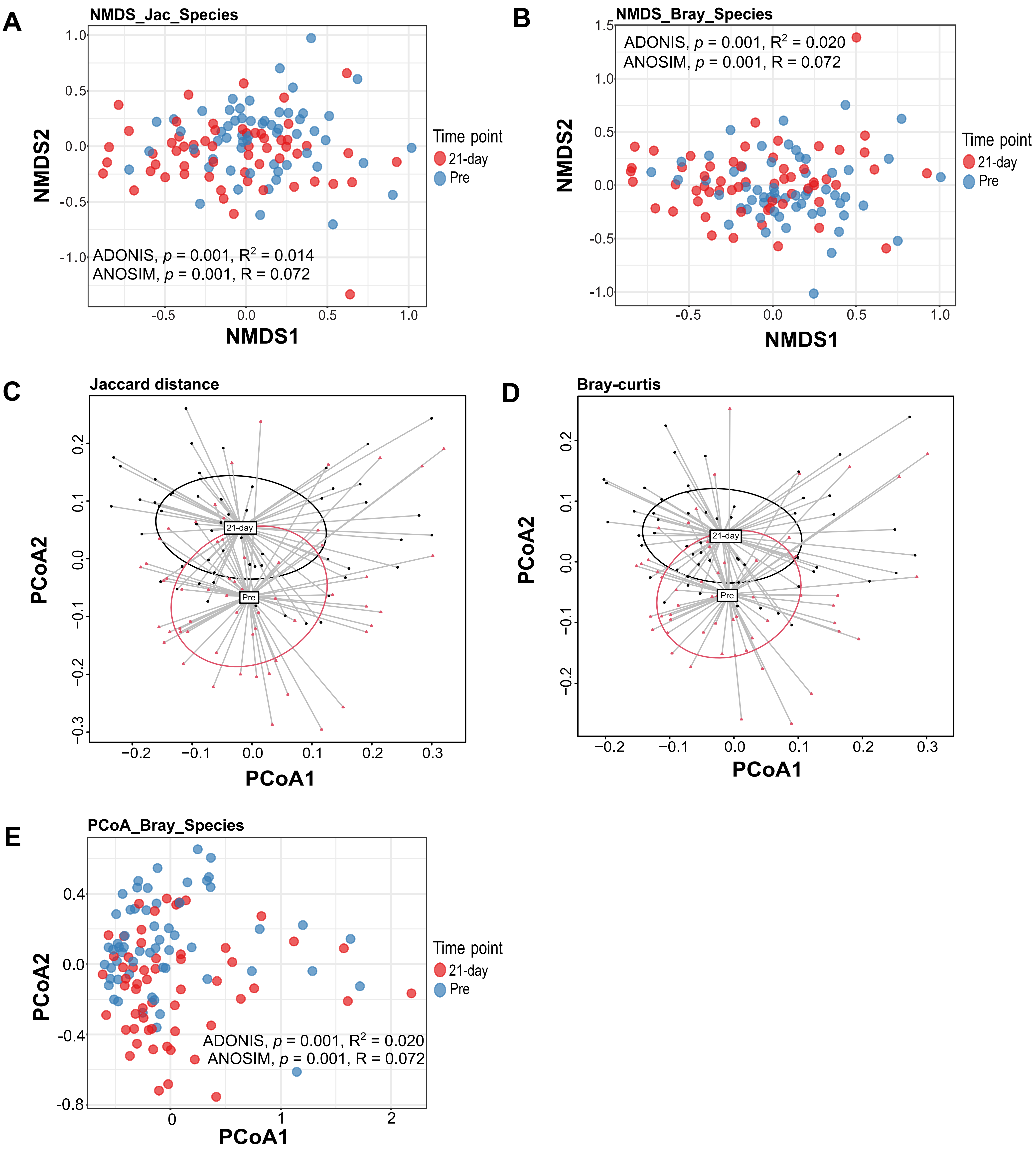

**Fig. S14**

Beta diversity comparing the gut microbiome between Pre-anthelminthic (Blue) and Post-anthelminthic (Red) among the Responders, visualized using Non-metric multi-dimensional scaling (NMDS) plot of **A** Jaccard distance (ADONIS: p=0.001, $R^2$=0.014; ANOSIM: p=0.001, R=0.072) and **B** Bray-curtis (ADONIS: p=0.001, $R^2$=0.020; ANOSIM: p=0.001, R=0.072). Principal Coordinates Analysis (PCoA) coordinate plot showing the beta-dispersion of **C** Jaccard distance and **D** Bray Curtis distance based on gut microbiota profile of the Responders, with Pre-anthelminthic (Red) and Post-anthelminthic (Black). **E** PCoA plot showing the beta diversity of Bray-curtis distances based on gut microbiota profile of Responders.

### Fig. S15

#### Uninfected

A
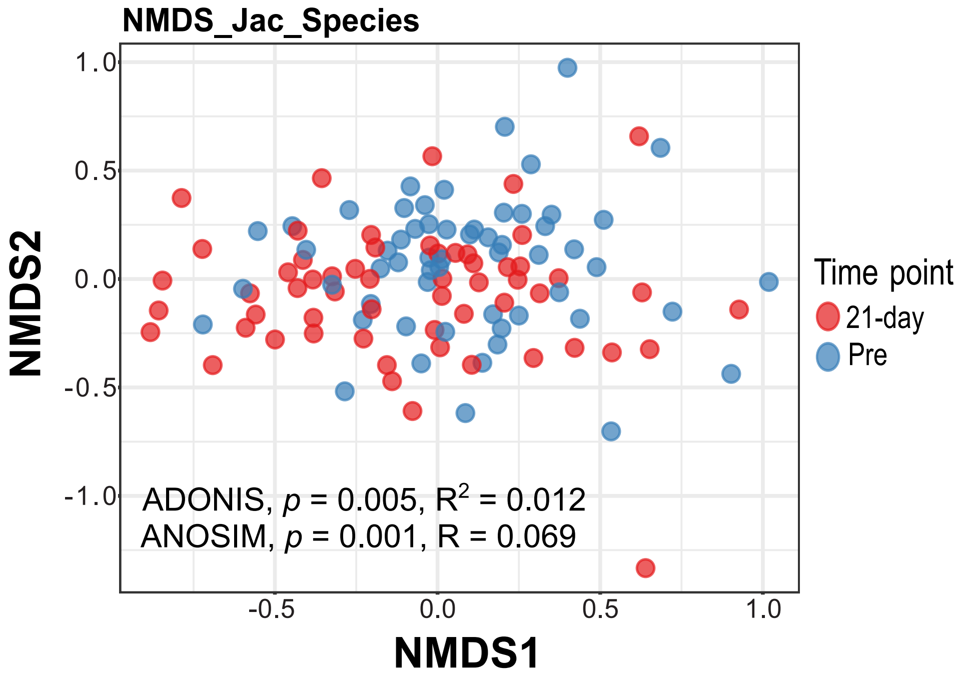


B
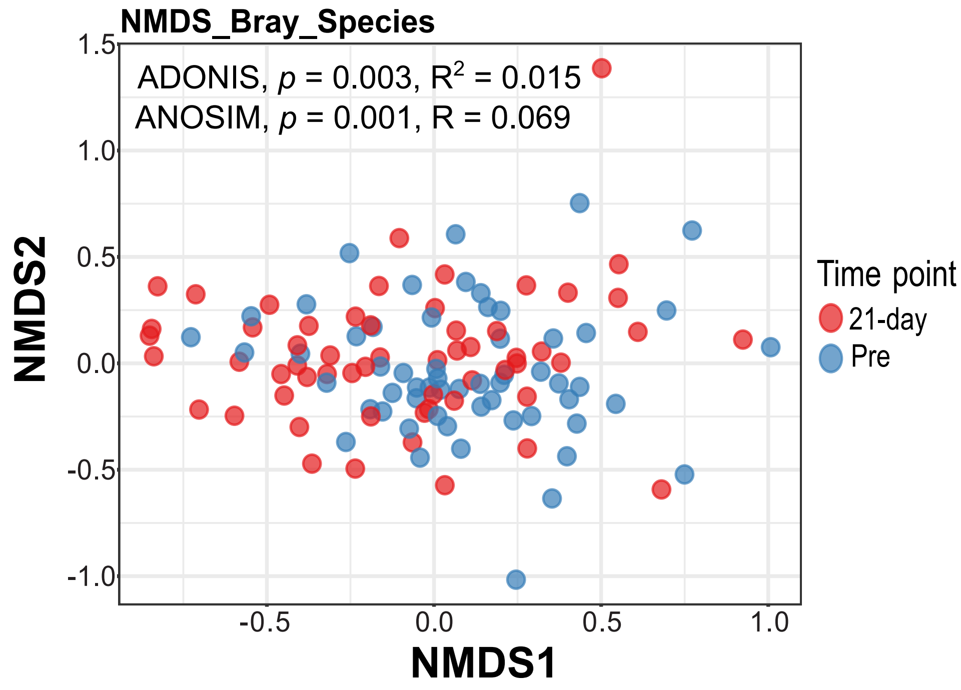


C
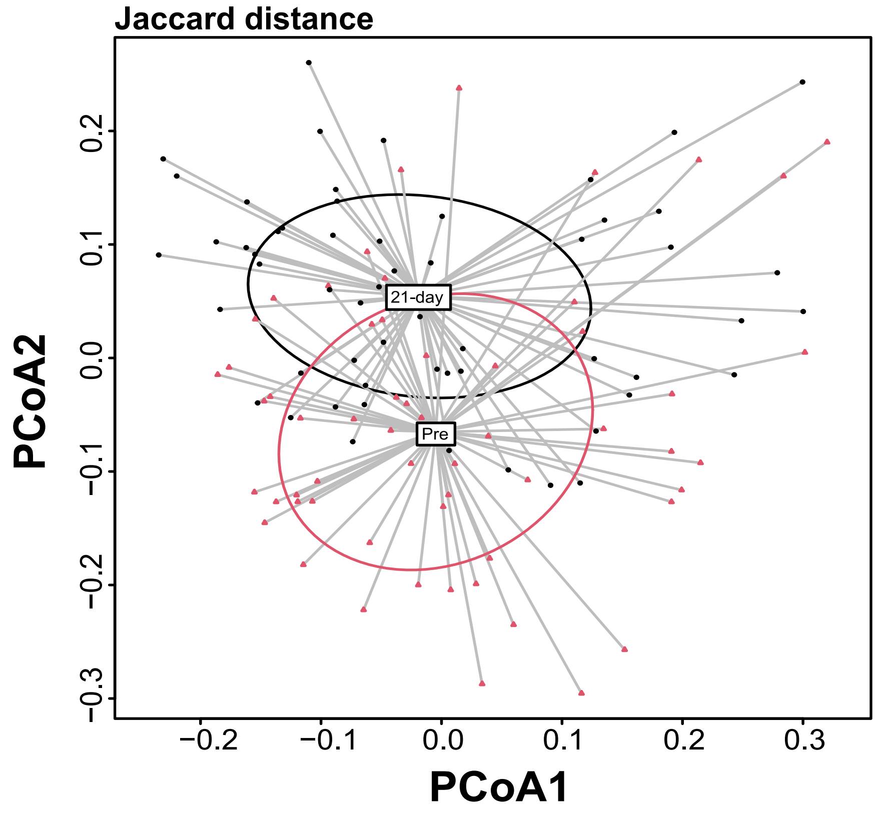


D
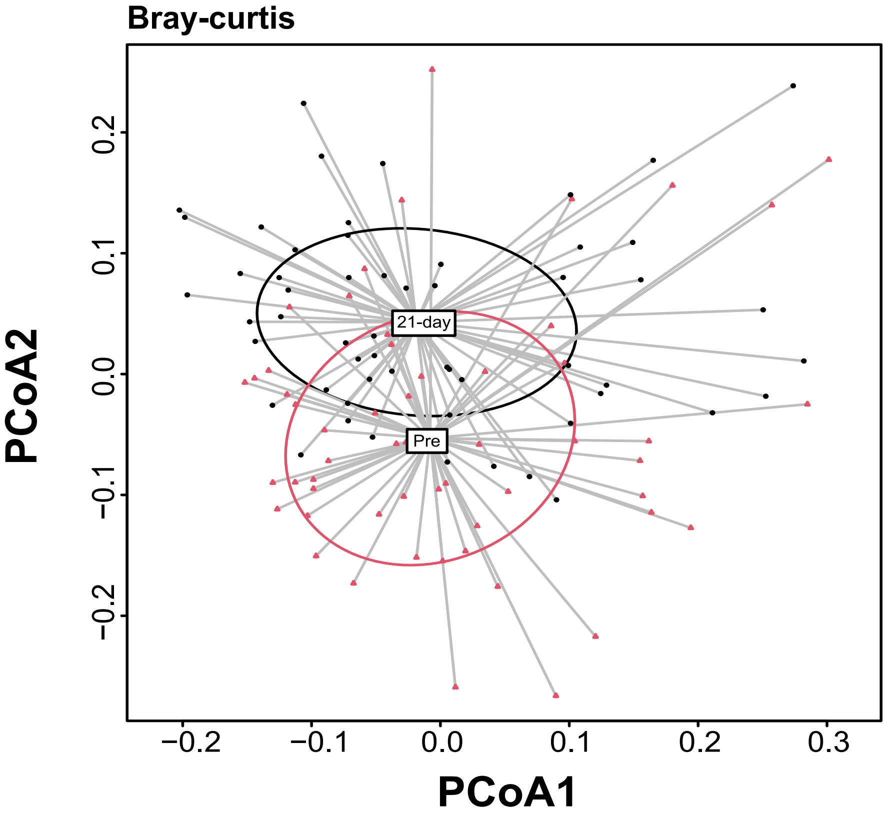


E
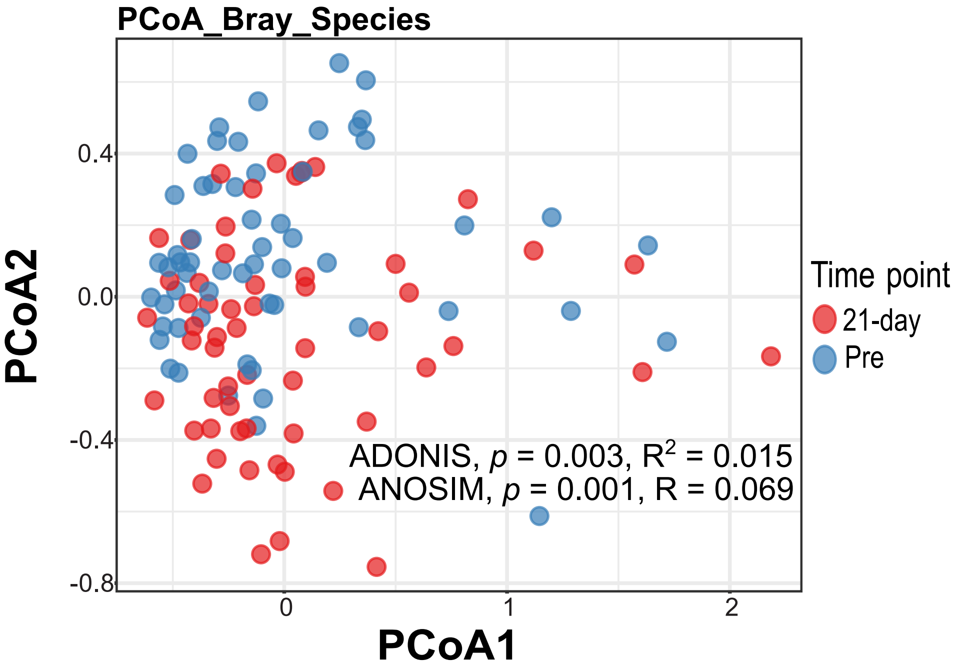

**Fig. S15**

Beta diversity comparing the gut microbiome between Pre-anthelminthic (Blue) and Post-anthelminthic (Red) among the Uninfected, visualized using Non-metric multi-dimensional scaling (NMDS) plot of **A** Jaccard distance (ADONIS: p=0.005, $R^2$=0.012; ANOSIM: p=0.001, R=0.069) and **B** Bray-curtis (ADONIS: p=0.003, $R^2$=0.015; ANOSIM: p=0.001, R=0.069). Principal Coordinates Analysis (PCoA) coordinate plot showing the beta-dispersion of **C** Jaccard distance and **D** Bray Curtis distance based on gut microbiota profile of the Uninfected, with Pre-anthelminthic (Red) and Post-anthelminthic (Black). **E** PCoA plot showing the beta diversity of Bray-curtis distances based on gut microbiota profile of the Uninfected.

Fig. S16

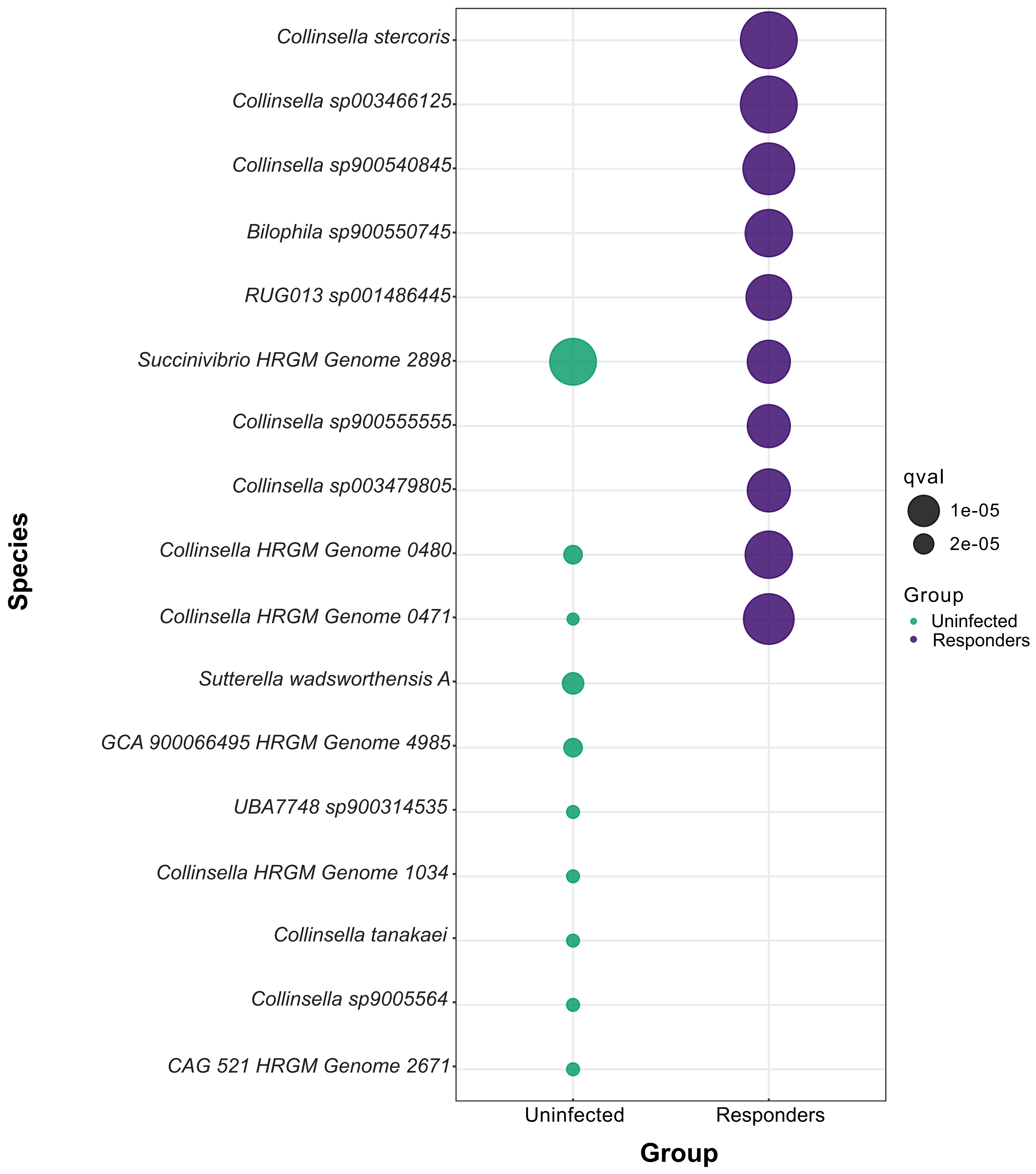

Species
Collinsella stercoris
Collinsella sp003466125
Collinsella sp900540845
Bilophila sp900550745
RUG013 sp001486445
Succinivibrio HRGM Genome 2898
Collinsella sp900555555
Collinsella sp003479805
Collinsella HRGM Genome 0480
Collinsella HRGM Genome 0471
Sutterella wadsworthensis A
GCA 900066495 HRGM Genome 4985
UBA7748 sp900314535
Collinsella HRGM Genome 1034
Collinsella tanakaei
Collinsella sp9005564
CAG 521 HRGM Genome 2671
Uninfected
Responders
Group
qval
1e-05
2e-05
Group
Uninfected
Responders

**Fig. S16**

Bubble plot shows the top 10 bacterial species that differentially abundant between Pre-anthelminthic and Post-anthelminthic  in Responders as well as the Uninfected based on the output of the Microbiome Multivariable Association with Linear Models 2 (MaAsLin2). The size of the bubble is negatively proportional to the p-value. The larger the bubble size displaying the lower p-value.

Fig. S17

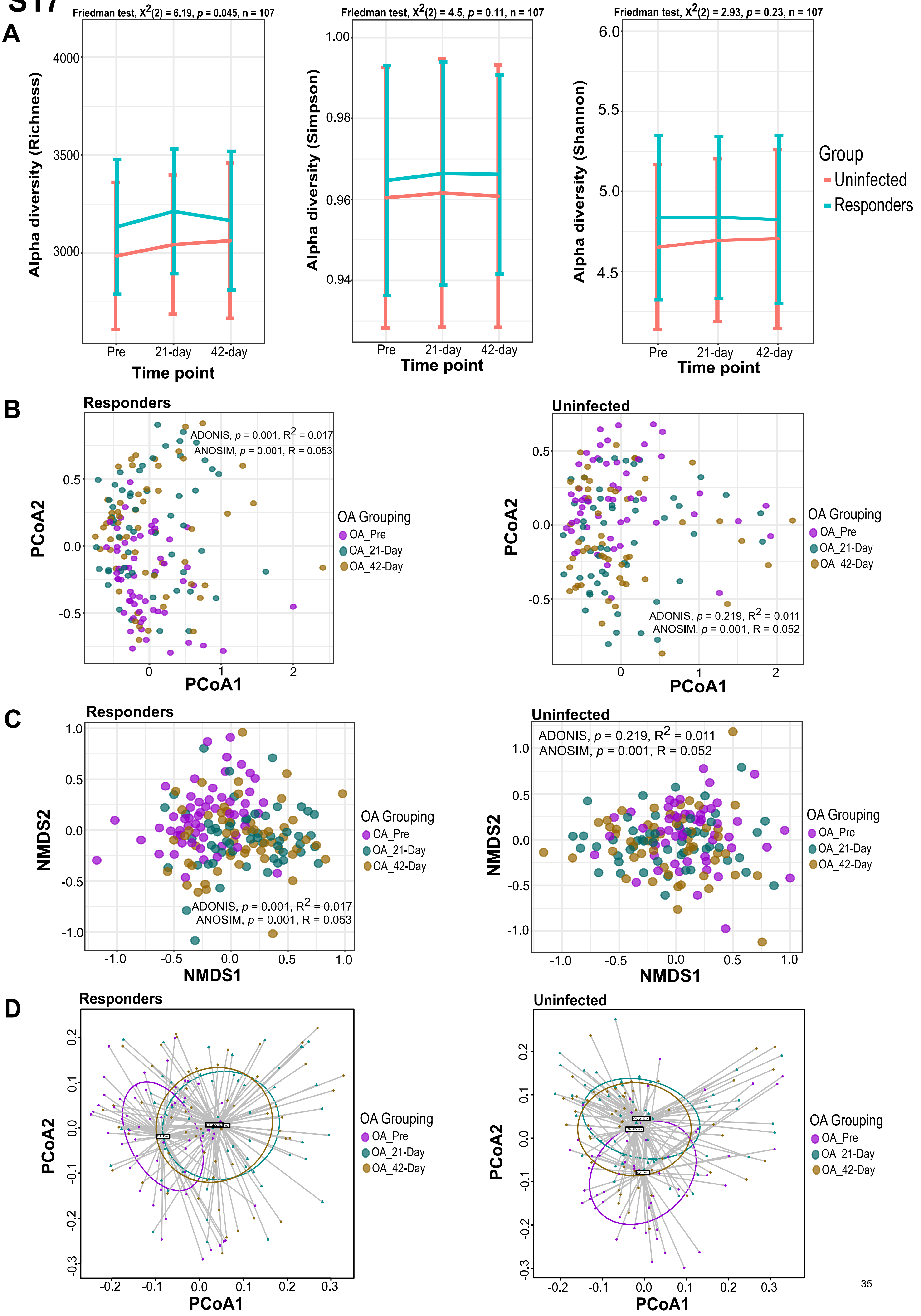
A
Friedman test, X2(2) = 6.19, p = 0.045, n = 107
Friedman test, X2(2) = 4.5, p = 0.11, n = 107
Friedman test, X2(2) = 2.93, p = 0.23, n = 107
Alpha diversity (Richness)
Alpha diversity (Simpson)
Alpha diversity (Shannon)
Time point
Pre
21-day
42-day
Group
Uninfected
Responders
B
Responders
Uninfected
ADONIS, p = 0.001, R2 = 0.017
ANOSIM, p = 0.001, R = 0.053
ADONIS, p = 0.219, R2 = 0.011
ANOSIM, p = 0.001, R = 0.052
PCoA1
PCoA2
OA Grouping
OA_Pre
OA_21-Day
OA_42-Day
C
NMDS1
NMDS2
D

**Fig. S17**

**A** Alpha diversity line plot showing the changes in the Richness, Shannon and Simpson diversity indices of the Orang Asli (OA) in Pre-anthelminthic, 21-Day and 42-Day Post-anthelminthic, with responders (Green) and uninfected (Red). The comparison of the alpha diversity index of all 3 timepoints were conducted using the Friedman test whereas between different two time points was tested using the Wilcoxon signed rank test. There are no statistical differences between groups. **B** Principal coordinates Analysis (PCoA), **C** Non-metric multi-dimensional scaling (NMDS), and **D** Beta-dispersion of Jaccard distance based on gut microbiota profile of the Pre-anthelminthic (Purple), 21-Day (Green) and 42-Day Post-anthelminthic (Gold) from the Responders (ADONIS: p=0.001, $R^2$=0.017; ANOSIM: p=0.001, R=0.053) (Left) and Uninfected (ADONIS: p=0.219, $R^2$=0.011; ANOSIM: p=0.001, R=0.052) (Right).

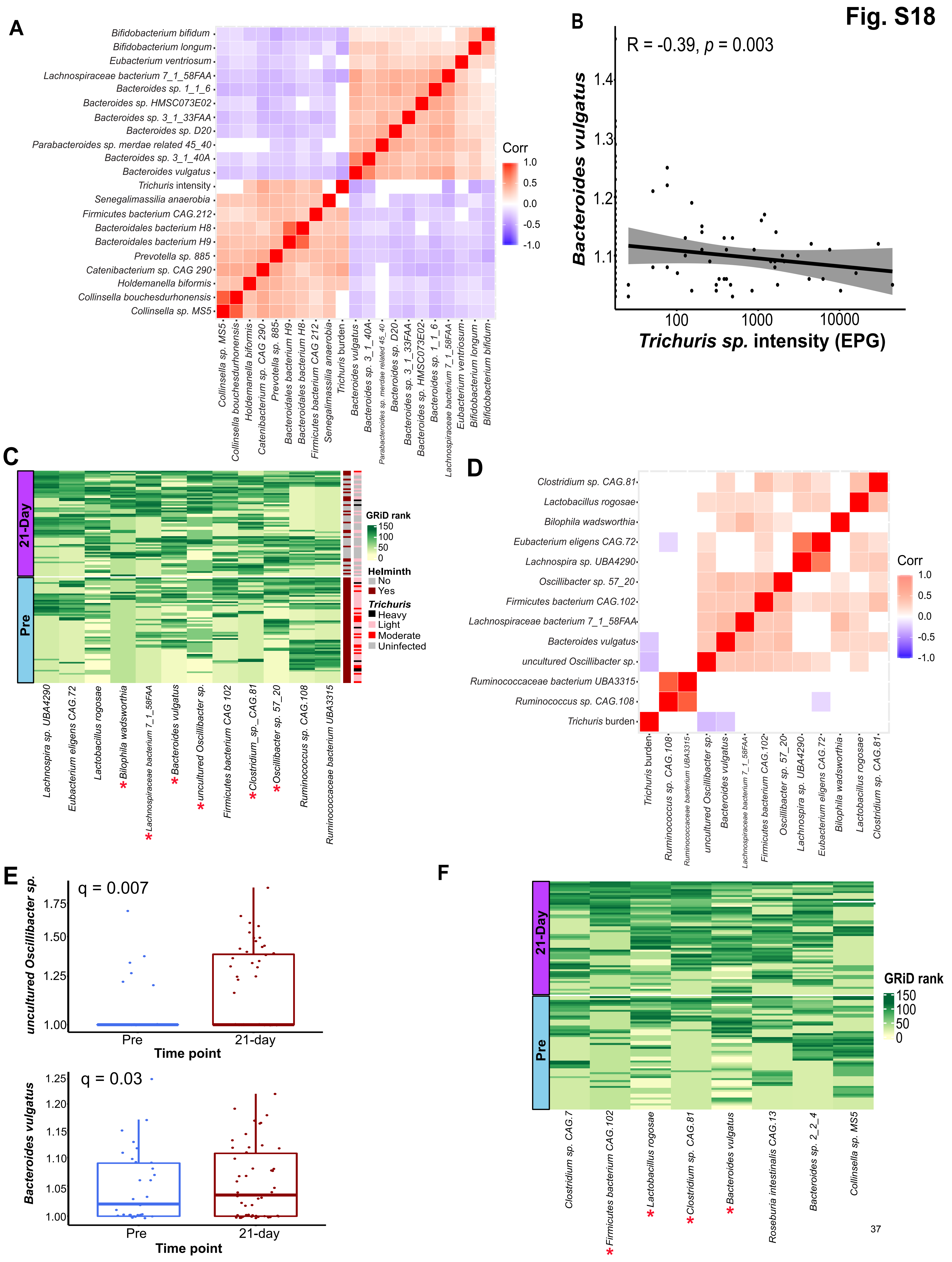

Fig. S18
A
B
R = -0.39, p = 0.003
Bacteroides vulgatus
Trichuris sp. intensity (EPG)
Corr
C
GRiD rank
Helminth
No
Yes
Trichuris
Heavy
Light
Moderate
Uninfected
21-Day
Pre
D
E
q = 0.007
uncultured Oscillibacter sp.
Time point
q = 0.03
Bacteroides vulgatus
Pre
21-day
F

**Fig. S18**

Growth Rate Index (GRiD) analysis of the gut microbial in the Orang Asli (OA) cohort. **A** Correlation matrix that showed to top 20 gut microbial species that correlate with the infection intensity of *Trichuris*. **B** Regression analysis between the abundance of *Bacteroides vulgatus* and the infection intensity of *Trichuris* (R=-0.39 and p=0.003). **C** Heatmap showing the replication the gut microbial species that are associated with intestinal helminth infection among the Responders. The first vertical side bar encodes the intestinal helminth infection while the second side bar indicates the infection intensity of the *Trichuris*. **D** Correlation matrix showing the Spearman correlation analysis between the top 12 gut microbial species and infection intensity of the *Trichuris* among the Responders. **E** Box plots showing the two bacteria [i.e., Uncultured *Oscilibacter* sp. (Top) and *Bacteroides vulgatus* (Bottom)] that are significantly negatively correlated with the infection intensity of the *Trichuris* in Responders. **F** Heatmap showing the replication of the gut microbial species that are associated with albendazole treatment among the Uninfected. Samples are shown in row by different time points (Pre-anthelminthic and Post-anthelminthic) whereas the rank of the GRiD score of each bacterium is shown in column.

### Fig. S19

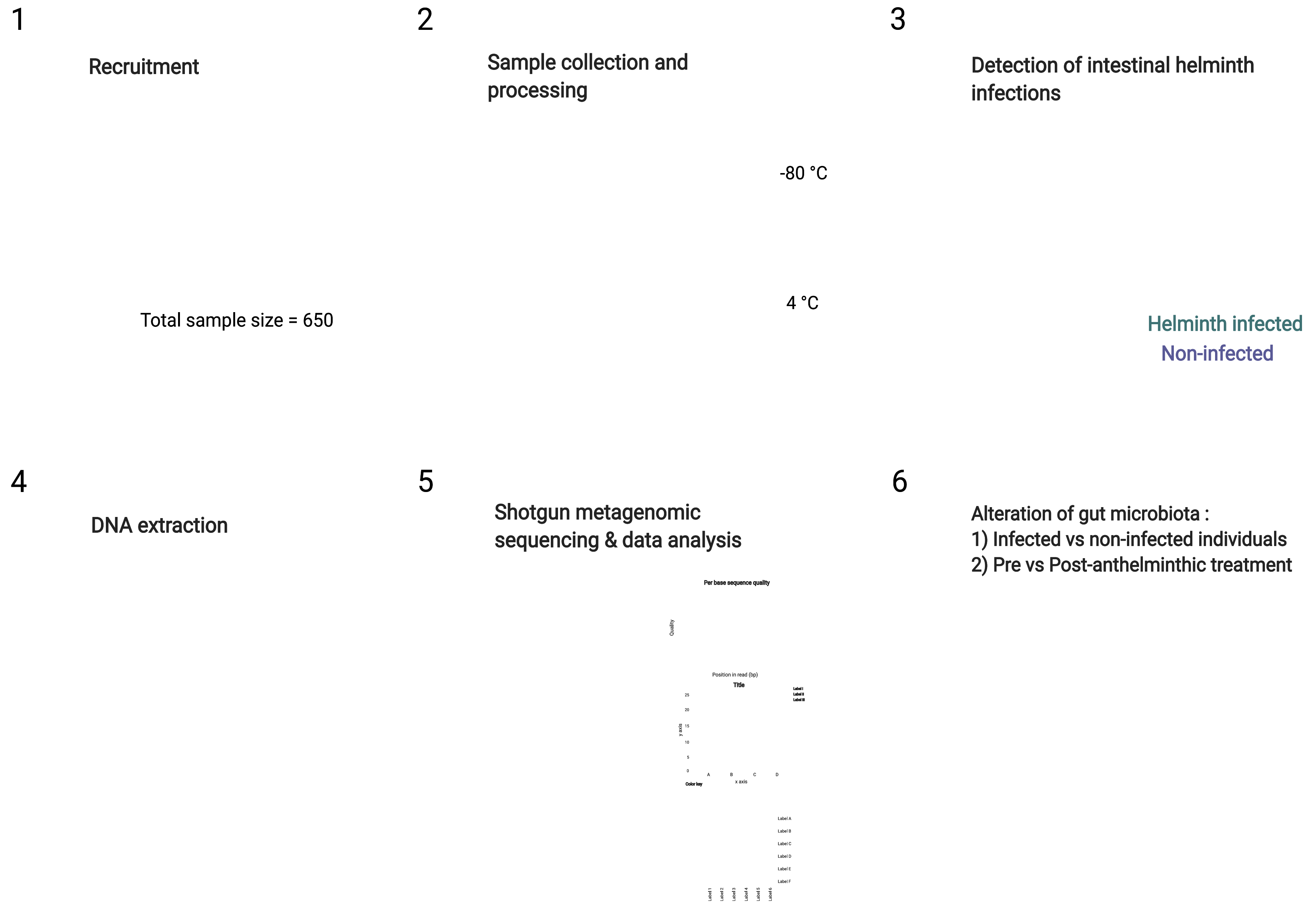


Created with BioRender.com

**Fig. S19**

The summary of the methodology from field work, sample collection, shotgun metagenomic sequencing and data analysis.

#### Fig. S20

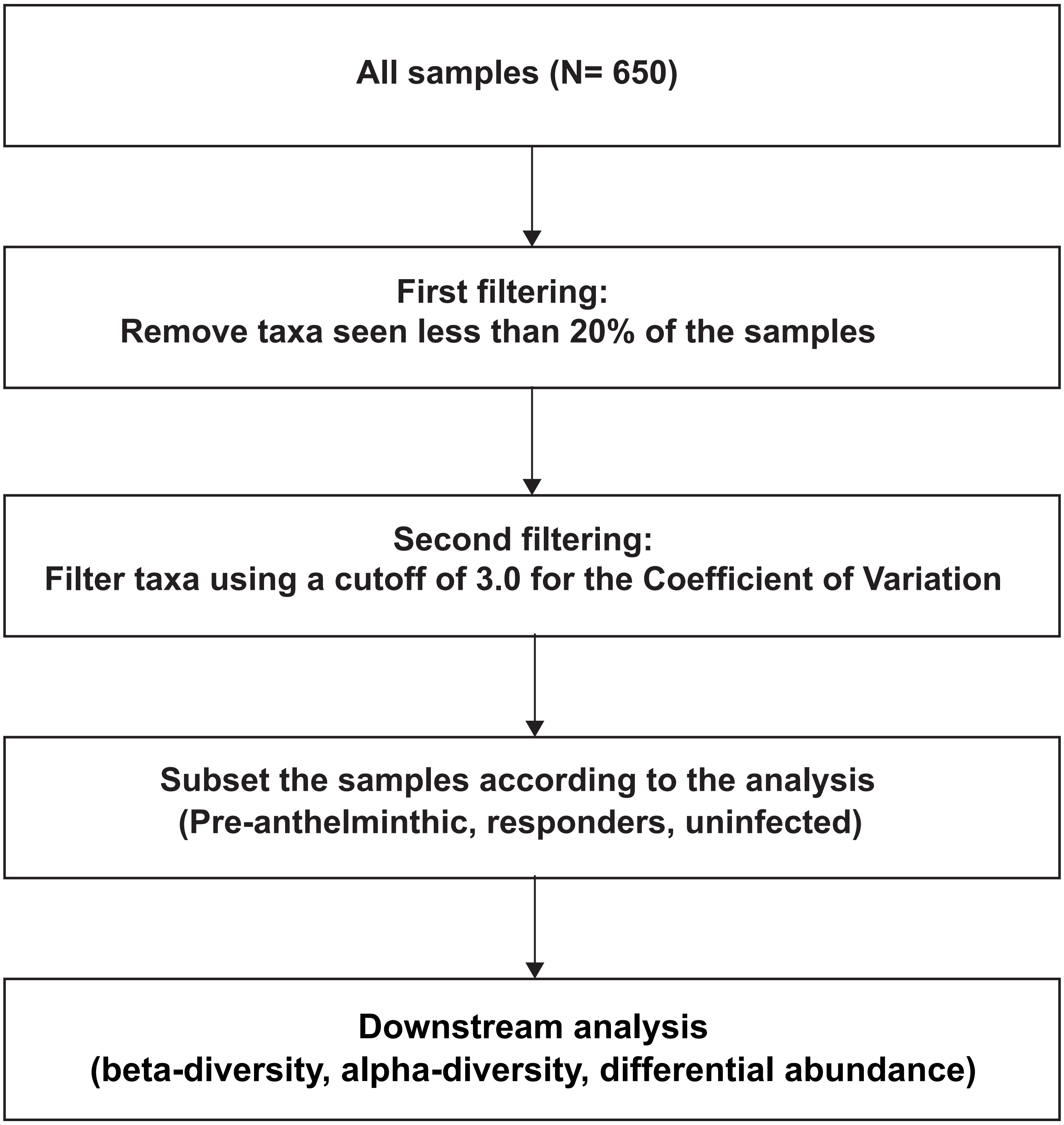
All samples (N= 650)
First filtering:
Remove taxa seen less than 20% of the samples
Second filtering:
Filter taxa using a cutoff of 3.0 for the Coefficient of Variation
Subset the samples according to the analysis
(Pre-anthelminthic, responders, uninfected)
Downstream analysis
(beta-diversity, alpha-diversity, differential abundance)

**Fig. S20**

A flow diagram showing the filtering steps before downstream analysis (beta-diversity, alpha diversity, and differential abundance).

Fig. S21

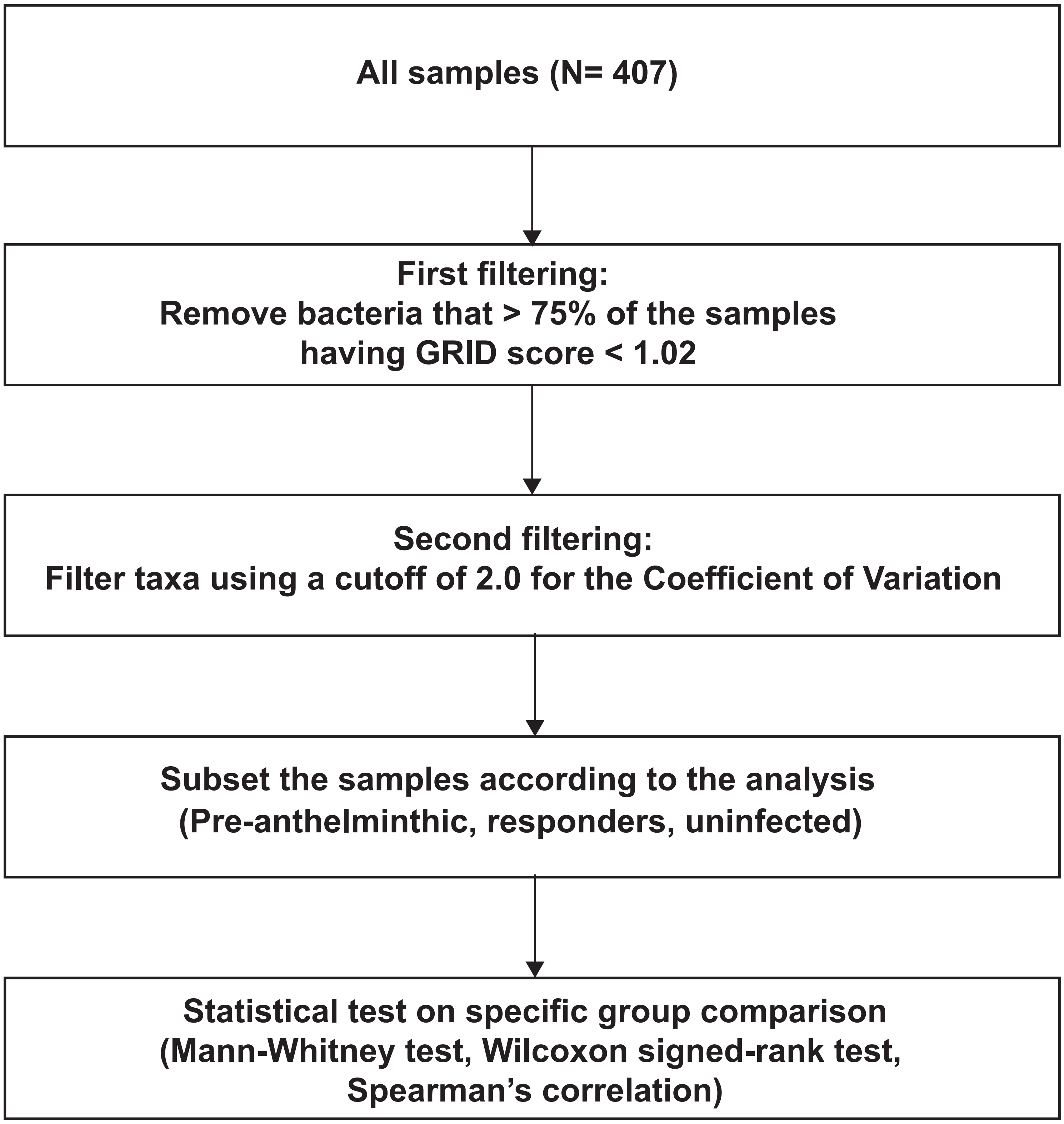
All samples (N= 407)
First filtering:
Remove bacteria that > 75% of the samples
having GRID score < 1.02
Second filtering:
Filter taxa using a cutoff of 2.0 for the Coefficient of Variation
Subset the samples according to the analysis
(Pre-anthelminthic, responders, uninfected)
Statistical test on specific group comparison
(Mann-Whitney test, Wilcoxon signed-rank test,
Spearman's correlation)

**Fig. S21**

Methodology for the evaluation of microbial growth rate in relation to helminth infection status in both cross-sectional and longitudinal phase using GRiD analysis.

Table S1: Relative impact of village, helminth infection and *Trichuris* infection on gut microbiome dissimilarity across samples (ADONIS, ANOSIM and Betadisper, permutation=999) in the cross-sectional analysis.

| Group | Jaccard | | | | | | | Bray-curtis | | | | | | | Sample size |
|---|---|---|---|---|---|---|---|---|---|---|---|---|---|---|---|
| | ADONIS | | | ANOSIM | | Tukey Test | | ADONIS | | | ANOSIM | | Tukey Test | | |
| | P value | F statistic | $R^2$ value | P value | R value | Compare group | P value | P value | F statistic | $R^2$ value | P value | R value | Compare group | P value | |
| Villages | 0.001 | 10.133 | 0.073 | 0.001 | 0.215 | Judah-Bangkong | 0.9999 | 0.001 | 15.058 | 0.105 | 0.001 | 0.215 | Judah-Bangkong | 0.9988 | All samples (n=650) |
| | | | | | | KL-Bangkong | 0.7904 | | | | | | KL-Bangkong | 0.7950 | |
| | | | | | | Legong-Bangkong | 0.0013 | | | | | | Legong-Bangkong | 0.0019 | |
| | | | | | | Rasau-Bangkong | 0.0018 | | | | | | Rasau-Bangkong | 0.0037 | |
| | | | | | | Sepat-Bangkong | 0.3373 | | | | | | Sepat-Bangkong | 0.3428 | |
| | | | | | | KL-Judah | 0.7525 | | | | | | KL-Judah | 0.8407 | |
| | | | | | | Legong-Judah | 0.0000 | | | | | | Legong-Judah | 0.0001 | |
| | | | | | | Rasau-Judah | 0.0001 | | | | | | Rasau-Judah | 0.0005 | |
| | | | | | | Sepat-Judah | 0.2062 | | | | | | Sepat-Judah | 0.2920 | |
| | | | | | | Legong-KL | 0.2259 | | | | | | Legong-KL | 0.2653 | |
| | | | | | | Rasau-KL | 0.2055 | | | | | | Rasau-KL | 0.2843 | |
| | | | | | | Sepat-KL | 0.9913 | | | | | | Sepat-KL | 0.9914 | |
| | | | | | | Rasau-Legong | 0.9999 | | | | | | Rasau-Legong | 1.0000 | |
| | | | | | | Sepat-Legong | 0.5371 | | | | | | Sepat-Legong | 0.5960 | |
| | | | | | | Sepat-Rasau | 0.4852 | | | | | | Sepat-Rasau | 0.6026 | |
| Helminth infection | 0.001 | 8.485 | 0.024 | 0.001 | 0.145 | Yes-No | 0.9475 | 0.001 | 12.783 | 0.035 | 0.001 | 0.145 | Yes-No | 0.9989 | All pre-treatment Orang Asli samples (n= 351) |
| *Trichuris* infection | 0.001 | 8.715 | 0.024 | 0.001 | 0.157 | Yes-No | 0.2250 | 0.001 | 13.099 | 0.036 | 0.001 | 0.157 | Yes-No | 0.2568 | All pre-treatmnet Orang Asli samples (n= 351) |

Table S2: Relative impact of deworming on gut microbiome dissimilarity of different group of Orang Asli samples based on their microbiome data in Pre and Post after the anthelminthic treatment (ADONIS, ANOSIM and Betadisper, permutation=999).

| Group | Jaccard | | | | | | | Bray-curtis | | | | | | | Sample size |
|---|---|---|---|---|---|---|---|---|---|---|---|---|---|---|---|
| | ADONIS | | | ANOSIM | | Tukey Test | | ADONIS | | | ANOSIM | | Tukey Test | | |
| | P value | F statistic | $R^2$ value | P value | R value | Compare group | P value | P value | F statistic | $R^2$ value | P value | R v alue | Compare group | P value | |
| Responders | 0.001 | 1.898 | 0.014 | 0.001 | 0.072 | Pre vs Post 1 | 0.9160 | 0.001 | 2.586 | 0.020 | 0.001 | 0.072 | Pre vs Post 1 | 0.9113 | Responders paired Orang Asli samples (n=66) |
| Uninfected | 0.006 | 1.382 | 0.012 | 0.001 | 0.069 | Pre vs Post 1 | 0.9182 | 0.003 | 1.766 | 0.015 | 0.001 | 0.069 | Pre vs Post 1 | 0.9024 | Uninfected paired Orang Asli samples (n=58) |

Table S3: Relative impact of the deworming on gut microbiome dissimilarity of different group of Orang Asli samples based on their gut microbiome data in Pre, 21-days and 42-days after the anthelminthic treatment (ADONIS, ANOSIM and Betadisper, permutation=999).

| Group | Jaccard | | | | | | | Bray-curtis | | | | | | | Sample size |
|---|---|---|---|---|---|---|---|---|---|---|---|---|---|---|---|
| | ADONIS | | | ANOSIM | | Tukey Test | | ADONIS | | | ANOSIM | | Tukey Test | | |
| | P value | F statistic | $R^2$ value | P value | R value | Compare group | P value | P value | F statistic | $R^2$ value | P value | R value | Compare group | P value | |
| Responders | 0.001 | 1.413 | 0.017 | 0.001 | 0.053 | Pre vs 21-Day<br>Pre vs 42-Day<br>21-Day vs 42-Day | 0.7096<br>0.9966<br>0.7567 | 0.001 | 1.808 | 0.021 | 0.001 | 0.053 | Pre vs 21-Day<br>Pre vs 42-Day<br>21-Day vs 42-Day | 0.6973<br>0.9974<br>0.7391 | Responders paired Orang Asli samples (n=51) |
| Uninfected | 0.219 | 1.067 | 0.014 | 0.002 | 0.037 | Pre vs 21-Day<br>Pre vs 42-Day<br>21-Day vs 42-Day | 0.9945<br>0.9902<br>0.9705 | 0.071 | 1.256 | 0.016 | 0.001 | 0.037 | Pre vs 21-Day<br>Pre vs 42-Day<br>21-Day vs 42-Day | 0.9916<br>0.9894<br>0.9632 | Uninfected paired Orang Asli samples (n=34) |
| All | 0.001 | 1.746 | 0.011 | 0.001 | 0.052 | Pre vs 21-Day<br>Pre vs 42-Day<br>21-Day vs 42-Day | 0.8404<br>0.9379<br>0.6399 | 0.001 | 2.412 | 0.015 | 0.001 | 0.052 | Pre vs 21-Day<br>Pre vs 42-Day<br>21-Day vs 42-Day | 0.8316<br>0.9269<br>0.6102 | All paired Orang Asli samples (n=85) |

Table S4. Heatmap shows the 61 bacterial species that significantly difference between intestinal helminth infected and uninfected subjects (Fig.4A)

| No | Species | P value | FDR |
|---|---|---|---|
| 1 | *Bacteroidales_bacterium_H9* | 5.57E-13 | 1.11E-11 |
| 2 | *Bacteroidales_bacterium_H8* | 9.88E-08 | 5.49E-07 |
| 3 | *Prevotella_sp._CAG.279* | 2.30E-07 | 1.09E-06 |
| 4 | *Clostridium_sp._CAG.127* | 0.0001 | 0.0003 |
| 5 | *Firmicutes_bacterium_CAG.212* | 5.43E-12 | 9.05E-11 |
| 6 | *Roseburia_sp._CAG.471* | 1.05E-05 | 3.90E-05 |
| 7 | *Collinsella_sp._MS5* | 3.57E-09 | 2.55E-08 |
| 8 | *Collinsella_bouchesdurhonensis* | 1.05E-07 | 5.55E-07 |
| 9 | *Catenibacterium_sp._CAG.290* | 1.65E-16 | 1.32E-14 |
| 10 | *Holdemanella_biformis* | 1.05E-08 | 6.55E-08 |
| 11 | *Prevotella_copri* | 2.40E-07 | 1.09E-06 |
| 12 | *Prevotella_sp._885* | 9.82E-10 | 8.93E-09 |
| 13 | *Firmicutes_bacterium_CAG.129_59_24* | 0.0073 | 0.0143 |
| 14 | *Oscillibacter_sp._ER4* | 3.73E-07 | 1.62E-06 |
| 15 | *Senegalimassilia_anaerobia* | 3.39E-14 | 8.48E-13 |
| 16 | *Oscillibacter_sp._57_20* | 0.01307 | 0.0233 |
| 17 | *Ruminococcus_sp._CAG.177* | 0.00827 | 0.0159 |
| 18 | *Faecalibacterium_sp._CAG.74_58_120* | 0.00491 | 0.0102 |
| 19 | *Firmicutes_bacterium_CAG.65_45_313* | 0.00128 | 0.0030 |
| 20 | *Eubacterium_sp._CAG.180* | 0.02108 | 0.0363 |
| 21 | *Eubacterium_sp._CAG.251* | 0.0122 | 0.0222 |
| 22 | *Faecalibacterium_sp._CAG.82* | 0.00346 | 0.0075 |
| 23 | *uncultured_Faecalibacterium_sp.* | 5.26E-05 | 0.0002 |
| 24 | *Faecalibacterium_prausnitzii* | 1.29E-05 | 4.60E-05 |
| 25 | *Clostridium_sp._CAG.265* | 1.21E-06 | 4.83E-06 |

| | | | |
|---|---|---|---|
| 26 | *Clostridium_sp._29_15* | 7.34E-05 | 0.0002 |
| 27 | *Eubacterium_ramulus* | 0.0002 | 0.0005 |
| 28 | *Ruminococcus_lactaris* | 0.0017 | 0.0039 |
| 29 | *Blautia_sp._CAG.237* | 0.0001 | 0.0004 |
| 30 | *Blautia_sp._KLE_1732* | 0.0057 | 0.0115 |
| 31 | *Blautia_obeum* | 0.0165 | 0.0290 |
| 32 | *Blautia_sp._SG.772* | 0.0049 | 0.0102 |
| 33 | *Eubacterium_sp._38_16* | 0.0119 | 0.0220 |
| 34 | *Bacteroides_sp._1_1_6* | 3.07E-11 | 3.83E-10 |
| 35 | *Bacteroides_sp._HMSC073E02* | 4.77E-08 | 2.81E-07 |
| 36 | *Bacteroides_sp._2_2_4* | 9.32E-07 | 3.88E-06 |
| 37 | *Parabacteroides_sp._merdae.related_45_40* | 1.86E-09 | 1.43E-08 |
| 38 | *Bacteroides_sp._D20* | 5.27E-10 | 5.85E-09 |
| 39 | *Parabacteroides_sp._20_3* | 1.57E-05 | 5.41E-05 |
| 40 | *Bacteroides_sp._3_1_33FAA* | 7.45E-12 | 1.06E-10 |
| 41 | *Bilophila_wadsworthia* | 3.42E-05 | 0.0001 |
| 42 | *Lachnospiraceae_bacterium_7_1_58FAA* | 1.51E-15 | 5.04E-14 |
| 43 | *Clostridium_sp._CAG.7* | 3.43E-06 | 1.32E-05 |
| 44 | *Clostridium_sp._CAG.81* | 0.0012 | 0.0029 |
| 45 | *Bacteroides_vulgatus* | 2.65E-16 | 1.32E-14 |
| 46 | *Bacteroides_sp._3_1_40A* | 9.19E-10 | 8.93E-09 |
| 47 | *uncultured_Oscillibacter_sp.* | 0.0001 | 0.0004 |
| 48 | *Roseburia_sp._UNK.MGS.15* | 3.28E-05 | 0.00010567 |
| 49 | *Eubacterium_ventriosum* | 8.67E-09 | 5.78E-08 |
| 50 | *Roseburia_intestinalis_CAG.13* | 0.0012 | 0.0029 |
| 51 | *Ruminococcus_bicirculans* | 0.0226 | 0.0376 |
| 52 | *Bacteroides_uniformis* | 0.0220 | 0.0372 |
| 53 | *Streptococcus_sp._HMSC065C01* | 0.0017 | 0.0038 |

| | | | |
|---|---|---|---|
| 54 | *Streptococcus_sp._SR4* | 2.86E-05 | 9.52E-05 |
| 55 | *Klebsiella_sp._KGM.IMP216* | 0.0058 | 0.0115 |
| 56 | *Clostridiales_bacterium_41_21_two_genomes* | 0.0001 | 0.0003 |
| 57 | *Bifidobacterium_longum* | 1.31E-09 | 1.09E-08 |
| 58 | *Bifidobacterium_sp._12_1_47BFAA* | 0.0002 | 0.0005 |
| 59 | *Bifidobacterium_adolescentis* | 0.0096 | 0.0182 |
| 60 | *Bifidobacterium_bifidum* | 1.91E-07 | 9.56E-07 |
| 61 | *Romboutsia_timonensis* | 0.0258 | 0.0423 |
